# Pathway-selective mitophagy regulates retinal physiology and neurogenic transitions in Muller glia

**DOI:** 10.64898/2026.05.17.725741

**Authors:** Aidan Anderson, Paula Rudzinska, Emer Chang, Dulani Wimalachandra, Kaouthar Bouzinab, Nada Alfahad, Samuel O Lord, Yu-Chiang Lai, Saaeha Rauz, Tim M Curtis, Graham R Wallace, Jose R Hombrebueno

**Author notes:** Correspondence to Dr Jose R Hombrebueno, Department of Inflammation and Ageing, School of Infection, Inflammation and Immunology, College of Medicine and Health, University of Birmingham, Edgbaston, Birmingham, B15 2TT, UK.

## Abstract

Mitochondrial quality control (MQC) is essential for retinal homeostasis, yet how distinct mitophagy pathways are coordinated within specialized retinal cell types remains poorly understood. Here, we show that Müller glia engage distinct mitophagy programmes that are differentially activated across physiological, metabolic stress, and differentiation contexts. Using pathway-resolved analyses supported by mouse and human single-cell transcriptomic datasets, we demonstrate that PINK1-dependent and receptor-mediated mitophagy pathways coexist within Müller glia and exhibit distinct functional and spatial regulation. To enable precise, time-resolved interrogation of these processes, we developed MQ-MG2, a spontaneously immortalised Muller glial model stably expressing the Mito-QC reporter while preserving endogenous mitophagy adaptors and metabolic features of primary Muller cells. Using this system, we identify context-dependent activation of mitophagy pathways with spatial relevance *in vivo* and reveal transient coordination of PINK1-dependent and receptor-associated mitophagy during Muller glial neurogenic differentiation. Suppression of fission-dependent mitophagy impaired the acquisition of complex neurite features in MQ-MG2, with a comparable phenotype observed following targeted PINK1 deletion in human neurogenic cells. Together, these findings position Muller glia as active integrators of mitochondrial quality control, capable of engaging distinct mitophagy programmes according to cellular context.

## INTRODUCTION

The retina is one of the most metabolically demanding tissues in the body, relyig on continuous, high-efficiency mitochondrial activity to support visual function. Maintenaning mitochondrial integrity is therefore critical for retinal homeostasis, and disruption of mitochondrial quality control (MQC) has been increasingly linked to retinal degeneration and metabolic disease [1–3]. Within this context, mitophagy, the selective autophagic degradation of mitochondria, serves as a pivotal MQC mechanism enabling rapid clearance of damaged organelles and facilitating cellular metabolic adaptation [4].

Multiple mitophagy pathways operate in mammalian cells, including the PTEN-induced kinase 1 (PINK1)–dependent pathway and receptor-mediated mechanisms involving BNIP3 and NIX [4]. Mitophagy is tightly coordinated with mitochondrial biogenesis (*de novo* synthesis of mitochondria), a process primarily regulated by peroxisome proliferator-activated receptor gamma coactivator-1α (PGC-1α) and mitochondrial transcription factor A (TFAM), which governs mtDNA replication, transcription and repair [5]. Mitochondrial dynamics, controlled by fusion and fission, also play a critical role in mitophagy by segregating damaged mitochondrial units for selective autophagic removal [3, 6].

While these pathways have been extensively characterised, how they are engaged under physiological conditions within metabolically specialized neural tissues remains poorly understood. In the retina, dysregulation of MQC has been implicated in several major ocular diseases, including diabetic retinopathy, age-related macular degeneration, and glaucoma, where impaired mitochondrial turnover contributes to cellular metabolic stress [1–3, 7]. Beyond disease, mitophagy is essential for neuronal differentiation and maturation, including in retinal ganglion cells, where it supports a transient metabolic shift toward glycolysis that precedes mitochondrial biogenesis and the establishment of oxidative phosphorylation capacity [8].

Within the retina, Muller glia function as a central metabolic hub. These radial glial cells span the full thickness of the retina, support neuronal survival, regulate neurotransmitter recycling, and maintain ionic and water homeostasis [9, 10]. Importantly, Muller glia also retain latent neurogenic competence [9, 11, 12], positioning them as a physiologically relevant system for examining how mitophagy is regulated during neural lineage commitment and their potential in retinal regeneration.

We and others have demonstrated that mitophagy is spatially enriched within the outer retina, a region populated by photoreceptors and Muller glia [2, 3, 13]. Emerging evidence further indicates that Muller glia actively contribute to MQC not only intrinsically but also through intercellular mechanisms such as trans-mitophagy, whereby damaged neuronal mitochondria are transferred to glial cells for degradation [14]. However, mechanistic understanding of how MQC is regulated within Muller glia remains limited, largely due to the absence of physiologically relevant systems that allow pathway-resolved and time-resolved analysis under basal and pathological conditions. Many existing approaches rely on biochemical assays or exogenous pathway overexpression, which can limit insight into endogenous regulatory mechanisms and complicate interpretation in metabolically specialized tissues such as the retina.

Here, we investigate how mitophagy is coordinated within retinal Muller glia under basal, metabolic stress, and differentiation conditions. Using a spontaneously immortalised Muller glia model derived from Mito-QC mice (MQ-MG2), we demonstrate preservation of endogenous PINK1-dependent and receptor-mediated mitophagy pathways, enabling pathway-resolved interrogation of mitochondrial turnover without genetic overexpression. Through this approach, we characterise time-resolved MQC remodelling during glial-to-neuronal differentiation and provide evidence that fission-dependent and PINK1-linked mitophagy support neurite maturation across mammalian neurogenic systems.

## RESULTS

### 1. Establishment of the MQ-MG2 Muller glial model preserving endogenous mitophagy programs

The Mito-QC mouse enables reliable quantification of mitophagy through a tandem mCherry-GFP reporter targeted to mitochondria (Figure 1A) [3, 15]. *In vivo*, mitophagy predominantly occurs at the outer retina, a region enriched with rod and cone photoreceptors as well as Muller glial radial processes, where Mito-QC is strongly expressed as identified by Glutamine Synthase (GS) immunostaining (Figure 1B, arrows). Similarly, the reporter is expressed in primary Muller cultures derived from Mito-QC mice (Mito-QC PMCs), which exhibit a flattened, elongated morphology (Figure 1C). During the expansion of Mito-QC PMCs, a subpopulation of cells continued to proliferate beyond the typical senescence period (passages 4–5), suggesting spontaneous immortalisation. To then establish a monoclonal Mito-QC Muller glia cell line, clonal isolation was performed via serial dilution. Eight clones were successfully isolated and expanded for further analysis. Among these, Clone 2 (herein referred to as MQ-MG2) exhibited the closest morphological resemblance to PMCs and displayed the strongest fluorescence of the Mito-QC reporter (Figure 1D-E). Based on these characteristics, MQ-MG2 was selected for detailed characterisation.

**Figure 1.**
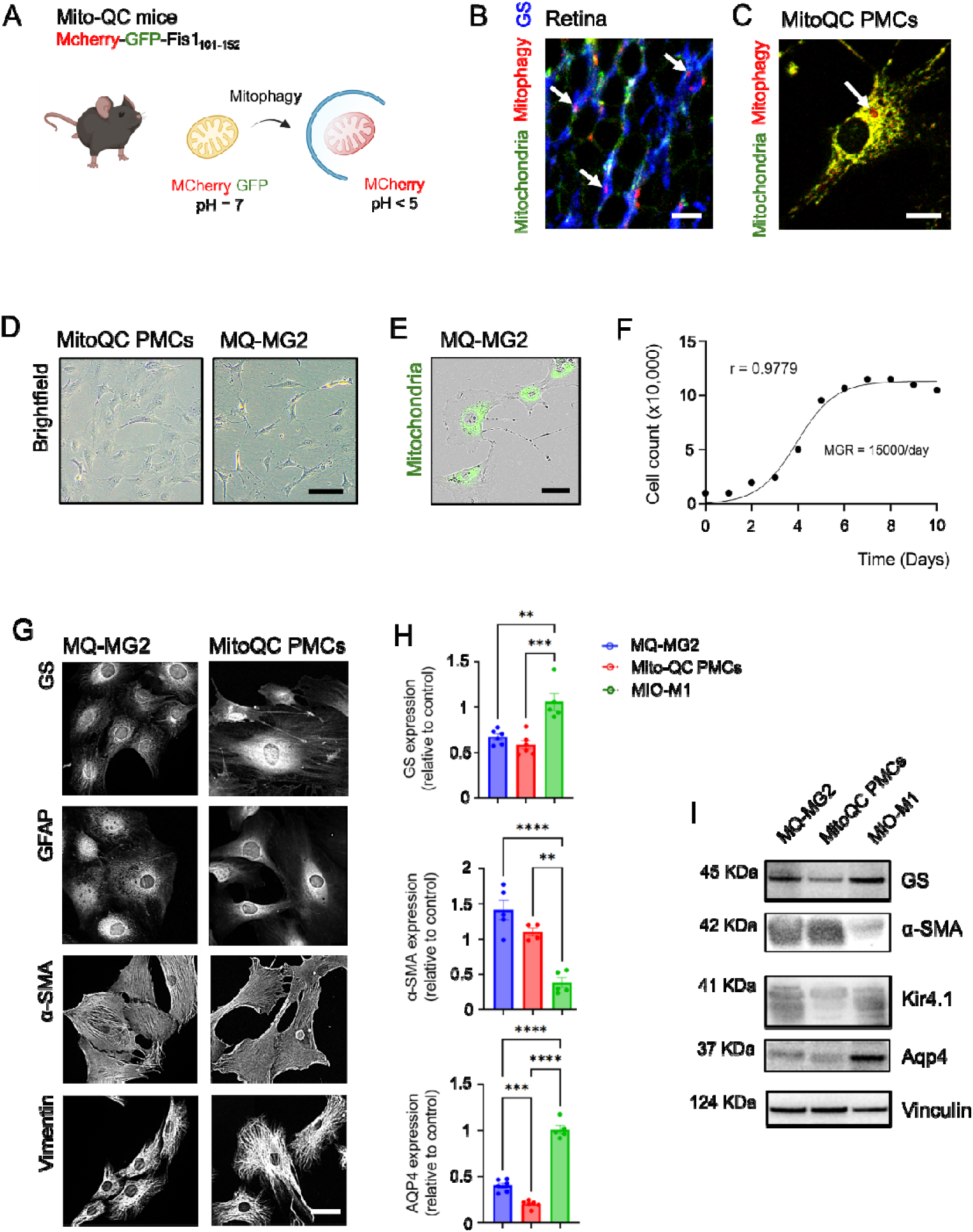
Development and characterization of immortalized MQ-MG2 Muller glial cells. **(A)** Assessment of mitophagy (mCherry-only puncta) and mitochondrial morphology (Fis1-GFP) using the Mito-QC reporter. **(B)** Retinal sections from 3-month-old Mito-QC reporter mice immunostained for glutamine synthase (GS) at the outer nuclear layer (ONL). A substantial proportion of mitolysosomes (arrows) localize within GS□ Muller glial processes. **(C)** Primary Muller cells from Mito-QC reporter mice (Mito-QC PMCs) showing mitochondrial networks and mitolysosomes (arrow). **(D-E)** Brightfield micrographs of Mito-QC PMCs and MQ-MG2 cells; panel (E) includes Mito-QC Fis1-GFP fluorescence using IncuCyte® Live-Cell. **(F)** Growth curve of MQ-MG2 cells over 10 days. **(G)** Representative micrographs of MQ-MG2 cells and Mito-QC PMCs stained for canonical Muller glial markers including GS, glial fibrillary acidic protein (GFAP), α-smooth muscle actin (α-SMA) and Vimentin. **(H-I)** Immunoblot analysis and quantification of GS, α-SMA, and Aquaporin-4 (AQP4) in lysates from MQ-MG2 cells, Mito-QC PMCs, and human MIO-M1 Muller cells. Protein levels were normalized to Vinculin. Data are presented as mean ± SD (*n* ≥ 4). **P < 0.01, ***P < 0.001, ****P < 0.0001. One-way ANOVA with Dunnett’s multiple comparison. Kir4.1, Inwardly Rectifying Potassium Channel 4.1; MGR, mean growth rate. Scale bars: 10 µm (B-C), 20 µm (G, E), 50 µm (D).

MQ-MG2 cells were cultured for over 50 passages while maintaining their proliferative capacity and stable expression of the Mito-QC reporter, as demonstrated by fluorescence microscopy and *Incucyte* live-cell imaging (Figure 1E). The doubling time of MQ-MG2 was 47.7 hours, with a mean growth rate of 15,000 cells/day, reaching a growth plateau by day 7 (Figure 1F). To confirm the Muller glia phenotype of MQ-MG2, we performed comprehensive immunocytochemistry and Western blot analyses using established Muller glia markers (Figure 1G-I), with mouse Mito-QC PMCs and the human cell line MIO-M1 as positive controls. MQ-MG2 cells broadly expressed canonical Muller glia markers, including GS, glial fibrillary acidic protein (GFAP), Vimentin, α-Smooth Muscle Actin (α-SMA), S100β, and Interleukin 33 (IL-33) (Figure 1G-I and Table S1). Additionally, MQ-MG2 expressed markers associated with specialized Muller glia functions, such as Aquaporin-4 and Kir4.1 (water homeostasis) and Heme-Oxygenase 1 (iron metabolism) (Figure 1G-I and Table S1). Western blot analysis confirmed that MQ-MG2 displayed expression levels of key markers similar to those in Mito-QC PMCs (Figure 1H-I). Notably, MQ-MG2 cells exhibited markedly higher levels of the proliferative marker Ki67 compared to Mito-QC PMCs, which instead showed features of cellular senescence, including reduced Lamin B1 expression (Supplemental Figure 1). These findings indicate that MQ-MG2 cells have undergone spontaneous immortalization while retaining the defining molecular characteristics of Muller glia.

### 2. Muller glia harbor PINK1-dependent and receptor-mediated mitophagy programs that can be pharmacologically engaged

To determine whether Muller glia possess the endogenous machinery required to support distinct mitophagy pathways, we first examined the expression of key mitophagy adaptors in healthy mouse and human retinas. Analysis of single-cell RNA sequencing datasets from the CZ CELLxGENE project [16, 17] revealed robust and cell-type–specific expression of both PINK1-dependent and receptor-mediated mitophagy components within Muller glia (Figure 2A-J). Specifically, we examined the expression of PINK1-dependent (*Pink1*, *Prkn*, *Phb2*, *Tbk1*, *Optn*) and receptor-mediated (*Bnip3*, *Nix*, *Fundc1*) mitophagy adaptors [4, 18]. Comparative analysis revealed that mouse and human retinas have a common expression pattern across main cellular subtypes, with *Nix* and *Prkn* being the most predominant transcripts, including in Muller glia (Figure 2A-J). Importantly, western blot analysis showed that MQ-MG2 cells express key mitophagy adaptors, including PINK1, PARKIN, BNIP3, and NIX at basal levels comparable to those observed in human MIO-M1 cells (Figure □2K). This suggests that Muller glia retain an inducible capacity for both PINK1-dependent and PINK1-independent mitophagy in the mammalian retina.

**Figure 2.**
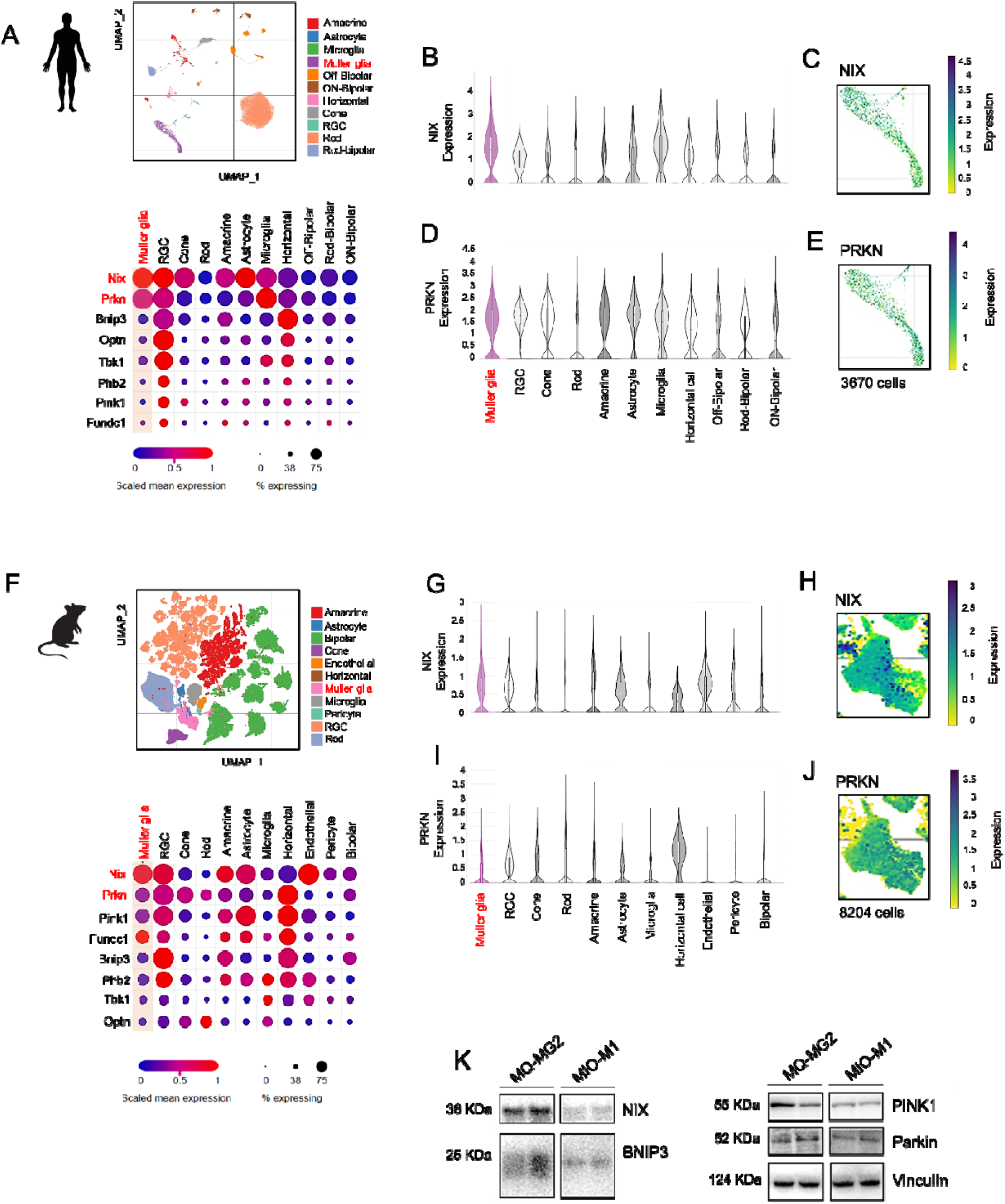
Human and mouse Muller glia express PINK1-dependent and receptor-mediated mitophagy adaptors that are retained in MQ-MG2 cells. **(A)** Uniform manifold approximation and projection (UMAP) of 51,645 human retinal cells profiled by scRNA-seq (CELLxGENE; Wang *et al.*, 2022) [16] derived from eight post-mortem retinas from four donors, resolving 11 major retinal cell types. Dot plot shows the percentage of expressing cells and scaled expression levels for transcripts associated with PINK1-dependent (*Pink1, Prkn, Phb2, Tbk1, Optn*) and receptor-mediated (*Bnip3, Nix, Fundc1*) mitophagy across retinal cell types. **(B-E)** Mean expression levels of *Nix* and *Prkn* across all human retinal cell types, with panels (C, E) highlighting expression within the Muller glia cluster (3,670 cells) on UMAP. **(F)** UMAP of 100,000 subsampled murine retinal cells compiled from seven curated public scRNA-seq datasets (CELLxGENE; Li *et al.*, 2024) [17], resolved into 11 major retinal cell types. Dot plot shows the percentage and scaled relative expression of PINK1-dependent and receptor-mediated mitophagy-related transcripts across the identified cell types. **(G-J)** Mean expression of *Nix* and *Prkn* in the mouse retina and within the Muller glia cluster, assessed as in (B-E). **(K)** Representative immunoblots showing endogenous expression of PINK1-dependent and receptor-mediated mitophagy adaptors in lysates from mouse MQ-MG2 cells and human MIO-M1 Muller cell lines. Vinculin was used as the loading control.

To characterise the inducible mitophagy capacity of Muller glia across distinct pathways, we first validated the responsiveness of MQ-MG2 to physiological stressors. Both hypoxia (1% O□) and amino acid starvation (HBSS) elicited a robust mitophagy response at 24 h, consistent with the stress-adaptive behaviour observed in primary Muller glia (Supplemental Figure 2A–C).

We then used MQ-MG2 to test whether mitophagy programmes inferred from scRNA-seq profiles could be selectively engaged by defined stimuli. To interrogate the PINK1-dependent axis, we applied established small-molecule modulators, including Kinetin and its potent derivate kinetin riboside (KR) [3, 19], as well as niclosamide [3, 20] and urolithin-A [3, 21]. Treatment with KR (0.7-5 µM), and niclosamide (1µM) effectively induced mitophagy in MQ-MG2, with a sensitivity comparable to that observed in Mito-QC PMCs (Figure 3A-C). The specificity of KR for activating PINK1-dependent mitophagy was validated at the molecular level by increased phospho-S65-ubiquitin (pS65-Ub) and PINK1 (Figure 4A–D). Moreover, KR reduced CISD1 levels (Figure 4E, H) independently of mitochondrial mass loss (Supplemental Figure 3A), consistent with ubiquitin-dependent turnover downstream of PINK1 pathway engagement [22, 23]. In contrast, expression of the receptor-mediated mitophagy adaptors BNIP3 and NIX remained unchanged (Figure 4E–G), supporting selective activation of the PINK1 pathway. Notably, neither kinetin nor urolithin-A induced mitophagy in MQ-MG2 or Mito-QC PMCs at any concentration tested (Figure 3A–C), consistent with our previous findings in Muller glia cultures [3].

**Figure 3.**
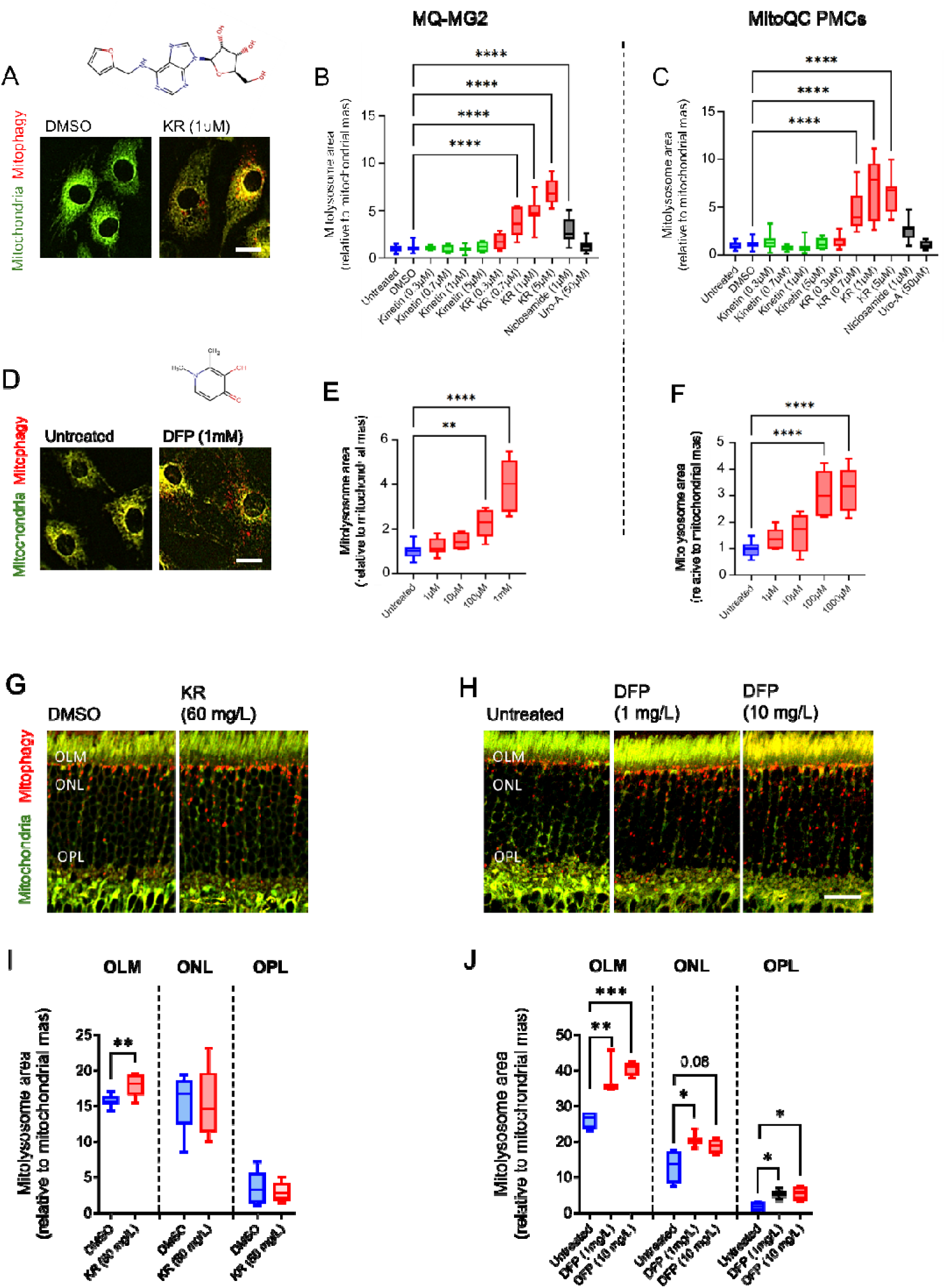
Pathway-selective activation of mitophagy is similarly inducible in Muller glia *in vitro* and in the outer retina *in vivo*. MQ-MG2 cells were treated with PINK1-dependent mitophagy activators (Kinetin, Kinetin Riboside [KR], Niclosamide, and Urolithin-A [Uro-A]) and a PINK1-independent enhancer (Deferiprone [DFP]). Responses were compared to primary Muller cells obtained from Mito-QC mice (Mito-QC PMCs). **(A, D)** Representative images of MQ-MG2 cells treated with KR (1□µM) or DFP (1□mM). **(B-C)** Quantification of mitophagy (mCherry-only puncta) in response to PINK1-dependent pharmacological enhancement. DMSO (0.1%) was used as vehicle control for all treatment groups **(E-F)** Quantification of mitophagy following treatment with the PINK1-independent enhancer DFP. **(G-J)** Fourteen-week-old Mito-QC mice were supplemented via drinking water with KR (60□mg/L), vehicle control (0.1% DMSO), or DFP (1□mg/L, 10□mg/L) for one week. **(G-H)** Representative retinal micrographs illustrating mitophagy levels across treatment groups in the outer retina. **(I-J)** Quantification of mitophagy in different subregions of the outer retina, including the outer limiting membrane (OLM), outer nuclear layer (ONL), and outer plexiform layer (OPL). Data are presented as box-and-whisker plots (median, interquartile range, and minimum/maximum values). (A-F) *n*□≥□4; (I) *n*□≥□7 eyes; (J) *n*□≥□3 eyes. *P < 0.05, **P < 0.01, ***P < 0.001, ****P < 0.0001. One-way ANOVA with Dunnett’s multiple comparison. (B–F, J); unpaired Student’s *t*-test (I). Scale bars: 10 µm (A, D), 20 µm (H).

**Figure 4.**
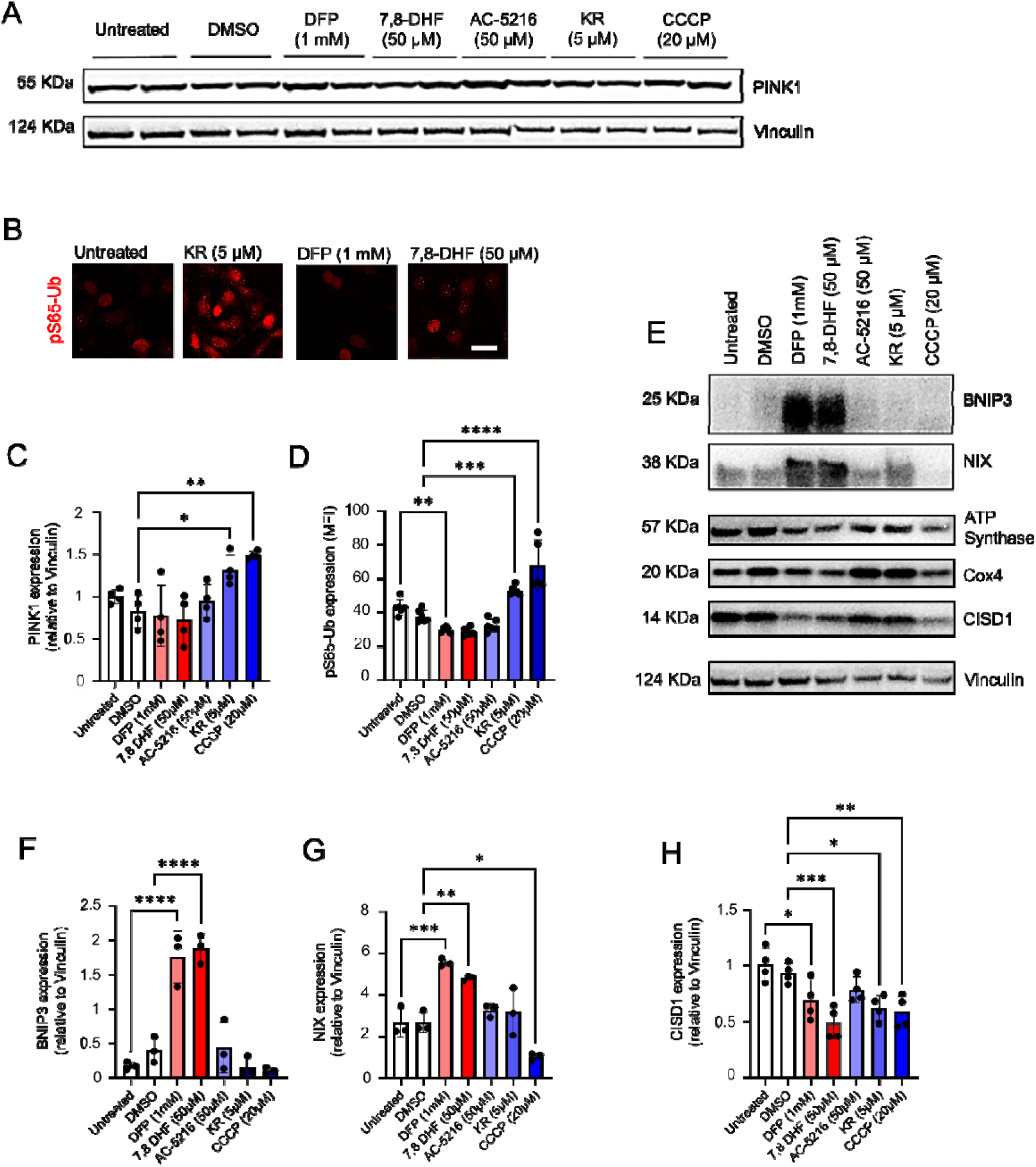
Pathway-specific activation of PINK1-dependent and receptor-mediated mitophagy in MQ-MG2 Muller glia. MQ-MG2 cells were treated with mitophagy activators—including Deferiprone (DFP), 7,8-Dihydroxyflavone (7,8-DHF), AC-516, Kinetin Riboside (KR), and Carbonyl cyanide m-chlorophenyl hydrazone (CCCP)—and pathway activation was assessed using biochemical and imaging approaches. **(A, E)** Representative immunoblots showing pathway activation following treatment with mitophagy activators. **(B)** Immunostaining for phosphorylated ubiquitin (pS65-Ub) as a readout of PINK1–Parkin–dependent mitophagy. **(C, F–H)** Immunoblot analysis and quantification of the mitophagy regulators PINK1, BNIP3, NIX, and CISD1 in MQ-MG2 cells across treatment conditions. Protein levels were normalized to Vinculin. **(D)** Quantification of pS65-Ub mean fluorescence intensity (MFI) across treatment groups. DMSO (0.1%) served as the vehicle control for all treatments except for DFP. Data are presented as mean□±□SD (*n*□≥□3). *P < 0.05, **P < 0.01, ***P < 0.001, ****P < 0.0001. One-way ANOVA with Dunnett’s multiple comparison. Scale bar: 20 µm.

To assess responsiveness to PINK1-independent mitophagy, cells were treated with the iron chelator deferiprone (DFP) [24, 25] or with 7,8-dihydroxyflavone (7,8-DHF) to activate the BDNF/TrkB pathway, which has been linked to BNIP3-associated mitophagy in specific cellular contexts [26]. Both DFP (Figure 3D-F) and 7,8-DHF (Supplemental Figure 2D-F) significantly induced mitophagy in MQ-MG2 and Mito-QC PMCs, accompanied by a net reduction in mitochondrial mass (Supplemental Figure 3B-C). Immunoblotting confirmed selective activation of PINK1-independent pathways, as treatment with DFP or 7,8-DHF increased BNIP3 and NIX expression (Figure 4E–G) without inducing pS65-Ub (Figure 4B, D).

In addition, we assessed the effects of translocator protein (TSPO) ligands, particularly AC-5216, based on prior evidence that TSPO regulates mitophagy and is expressed in retinal Muller glia under physiological and pathological conditions [27, 28]. Treatment with AC-5216 induced mitophagy in MQ-MG2 cells, with comparable responses in Mito-QC PMCs (Supplemental Figure 2G–I), without significant changes in mitochondrial mass (Supplemental Figure 3D). Immunoblotting revealed no detectable changes in PINK1-dependent markers, PINK1-independent adaptors, or CISD1 (Figure 4), indicating that mitophagy activation via TSPO occurs through an alternative pathway, consistent with previous reports [27].

To determine whether pathway-specific mitophagy modulation observed in Muller glia translates *in vivo*, Mito-QC mice were treated with KR or DFP administered via drinking water. Consistent with our previous observations in diabetes models, KR elicited mitophagy localised to the outer limiting membrane (OLM) (Figure 3G, I). Notably, KR did not result in a significant reduction in mitochondrial mass (p = 0.09; Supplemental Figure 4A), consistent with our *in vitro* findings suggesting compensatory coupling of PINK1-dependent mitophagy to mitochondrial biogenesis. In contrast, DFP enhanced mitophagy at lower concentrations across multiple outer retinal layers, including the OLM, outer nuclear layer (ONL), and outer plexiform layer (OPL) (Figure 3H, J), and was associated with a significant decrease in mitochondrial mass (Supplemental Figure 4B). This spatial pattern closely mirrored scRNA-seq profiles of the outer retina, which showed predominant expression of NIX compared to PINK1-dependent pathways (Figure 2).

Similarly, 7,8-DHF supplementation reduced mitochondrial mass in both Mito-QC mice (lower Fis1-GFP signal) and non-Mito-QC mice, as indicated by decreased Cox4 and TFAM levels (Supplementary Figure □4C–F). Together, these findings demonstrate that Muller glia harbor multiple inducible mitophagy programs, and that activation of receptor-associated BNIP3/NIX pathways is preferentially associated with net mitochondrial clearance.

### 3. Muller glia engage mitophagy to preserve mitochondrial function under stress

We next profiled mitochondrial respiration and glycolytic activity in MQ-MG2 cells using the Seahorse XF96e extracellular flux assay. Oxygen consumption rates (OCR) and proton efflux rates (PER) were measured under both basal and stressed conditions, using the mitochondrial uncoupler BAM15 in combination with monensin to assess maximum glycolytic capacity (Figure 5) [29, 30].

**Figure 5.**
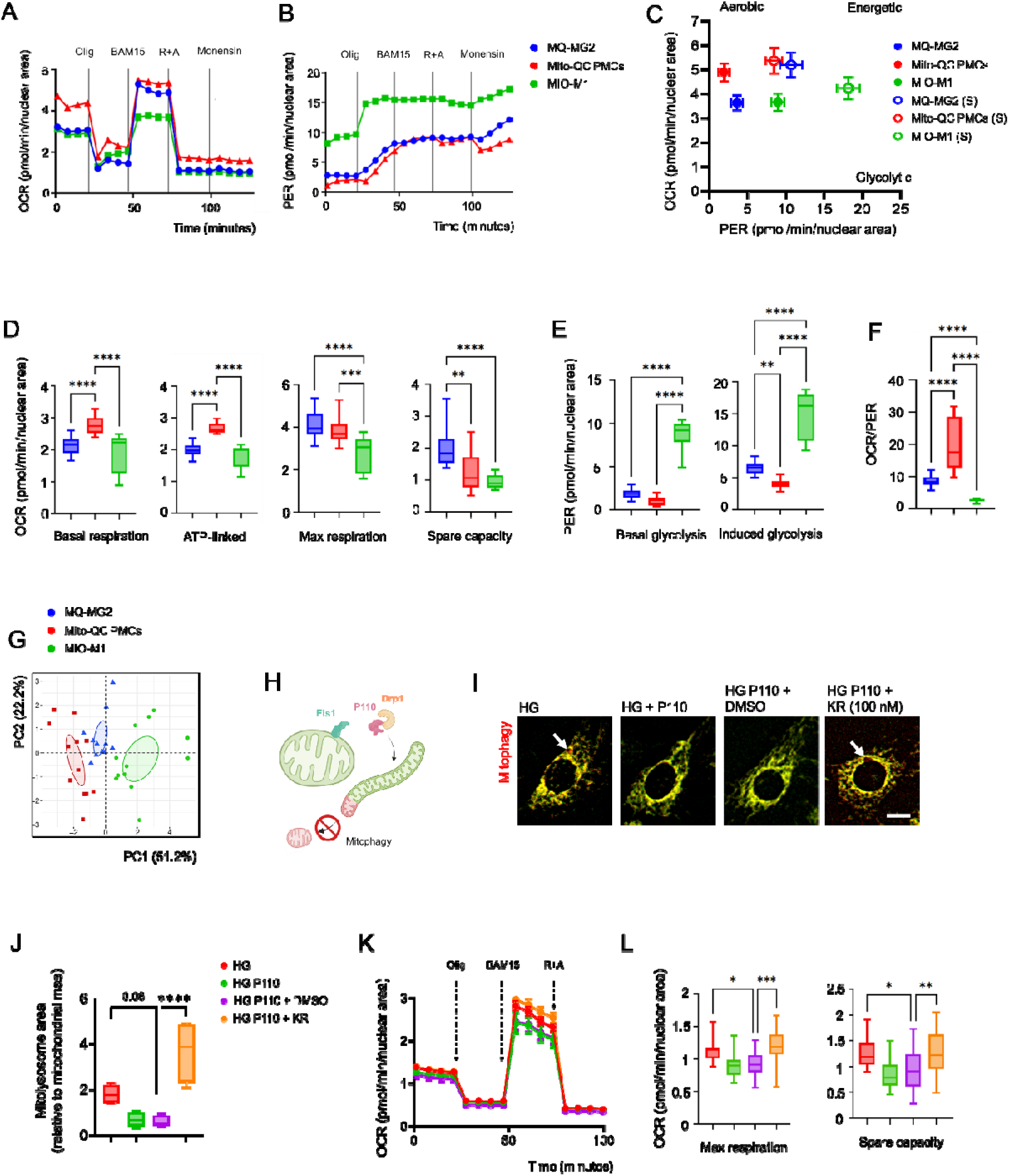
Mitophagy preserves mitochondrial function and bioenergetic capacity under metabolic stress in Muller glia. (A-B) Representative Seahorse assay of metabolic flux using *Cell Mito Stress Test* (left) and *Glyco Stress Test* (right) in MQ-MG2 cells, primary Muller cells from Mito-QC reporter mice (Mito-QC PMCs), and human MIO-M1 cells. **(C)** Bioenergetic phenotype maps showing oxygen consumption rate (OCR) versus proton efflux rate (PER) under basal and stressed conditions across all Muller glial cell types. **(D-E)** Quantification of OCR (mitochondrial respiration) and PER (glycolytic activity) under basal and stressed conditions. **(F)** The OCR:PER ratio under basal conditions, comparing the bioenergetic reliance of each Muller glial cell type. **(G)** Principal component analysis (PCA) with confidence ellipses visualizing clustering of Muller glial cell types based on OCR and PER parameters under basal and stressed conditions**. (H-L)** MQ-MG2 cells were cultured under hyperglycaemia for 3 days (HG; 30.5□mM) and mitochondrial fission antagonized with P110 to replicate diabetic stress, with or without pharmacological activation of PINK1 using Kinetin Riboside (KR; 100 nM) or vehicle control (0.1% DMSO). **(I-J)** Quantification of mitophagy (mCherry-only puncta, arrows) in different treatment groups. **(K-L)** Representative *Cell Mito Stress Test* traces and quantification of OCR parameters reflecting mitochondrial fitness, including maximal respiration and spare respiratory capacity. Data are presented as mean ± SE (A-C, K); or box-and-whisker plots (median, interquartile range, and minimum/maximum values) (D-F, J, L). *n*□≥□12 (A-F, K-L); *n*□=□4 (J). *P < 0.05, **P < 0.01, ***P < 0.001, ****P < 0.0001. One-way ANOVA with Dunnett’s multiple comparison. Olig, oligomycin; R+A, rotenone + antimycin. Scale bar: 5 µm.

Under basal conditions, both MQ-MG2 and MIO-M1 cells exhibited lower mitochondrial and ATP-linked respiration compared to PMCs (Figure 5A, D). However, upon BAM15-mediated mitochondrial uncoupling, MQ-MG2 reached maximum respiration levels comparable to PMCs (Figure 5A, D). Notably, MQ-MG2 demonstrated superior spare respiratory capacity compared to both PMCs and MIO-M1 cells, indicating a greater capacity for mitochondrial respiration under energy demand (Figure 5A, D). These findings were validated using the classical *Mito Stress test* with FCCP as an uncoupler, where MQ-MG2 exhibited maximal respiration similar to PMCs but with superior spare respiratory capacity (Supplemental Figure 5).

Basal glycolysis was similar between MQ-MG2 and PMCs but was elevated in MIO-M1 cultures (Figure 5B, E). Upon monensin treatment, glycolytic capacity increased significantly in MQ-MG2 compared to PMCs, though it remained lower than in MIO-M1 cultures (Figure 5B, E). Bioenergetic phenotype maps (OCR vs PER) showed that MQ-MG2 more closely resembled PMCs under both basal and stressed conditions, while MIO-M1 remained highly glycolytic (Figure 5C, F). This similarity was further supported by Principal Component Analysis (PCA), which showed MQ-MG2 clustering more closely with PMCs than with MIO-M1 cells (Figure 5G).

We previously reported that mitochondrial hyperfusion impairs turnover and leads to metabolic stress in advanced neurodegenerative diabetic retinopathy [3]. This pathology can be modelled in Mito-QC PMCs by inhibiting mitochondrial fission with the P110 peptide (a DRP1 specifiic inhibitor) under hyperglycaemic conditions [3]. To assess whether MQ-MG2 can model DR-associated mitochondrial pathology and its rescue via mitophagy induction, we treated MQ-MG2 cells with P110 (Figure 5H). P110 treatment inhibited mitophagy (Figure 5I-J), inducing bioenergetic stress under hyperglycaemic conditions, as indicated by reduced maximal and spare respiratory capacity (Figure 5K-L), mirroring the response we previously observed in Mito-QC PMCs [3]. Treatment with KR at low nanomolar concentrations (100 nM) rescued mitophagy and restored bioenergetic profiles in MQ-MG2 under diabetic hyperfusion conditions (Figure 5I-L), Together, these findings confirm that MQ-MG2 recapitulates mitochondrial phenotypes previously identified in diabetic retinopathy models, supporting bioenergetic and mechanistic analysis of stress-associated MQC adaptation.

### 4. Fission-dependent mitophagy contributes to mitochondrial remodelling during Muller glial neurogenic differentiation

Several immortalised Muller cell lines, including MIO-M1, have been reported to exhibit retinal stem cell properties, such as differentiation into retinal neuronal-like cells and/or neurosphere formation [31]. Given the glial origin of MQ-MG2, we assessed the expression of neural progenitor markers, using MIO-M1 cells as a positive control. Notably, MQ-MG2 expressed canonical retinal stem cell transcription factors, including Pax6, Sox2, and Notch1, along with detectable expression of early neuronal cytoskeletal markers, such as Nestin and βIII-tubulin (Figure 6A-B). Exposure of MQ-MG2 to neurogenic commercial media (see Methods) induced rapid neuronal-like differentiation, evident from the formation of neural projections within 1-3 hours (Figure 6C-E), as further captured by *Incucyte* live-cell imaging (Figure 6F). Over 2-4 days, MQ-MG2 cells developed extended neuronal-like processes and formed neurospheres (Figure 6C). These morphological changes were accompanied by a sharp upregulation of mature neuronal markers, including heavy chain neurofilament (NF-H) and β-III-tubulin (Figure 6D-E), confirming their neuronal differentiation potential.

**Figure 6.**
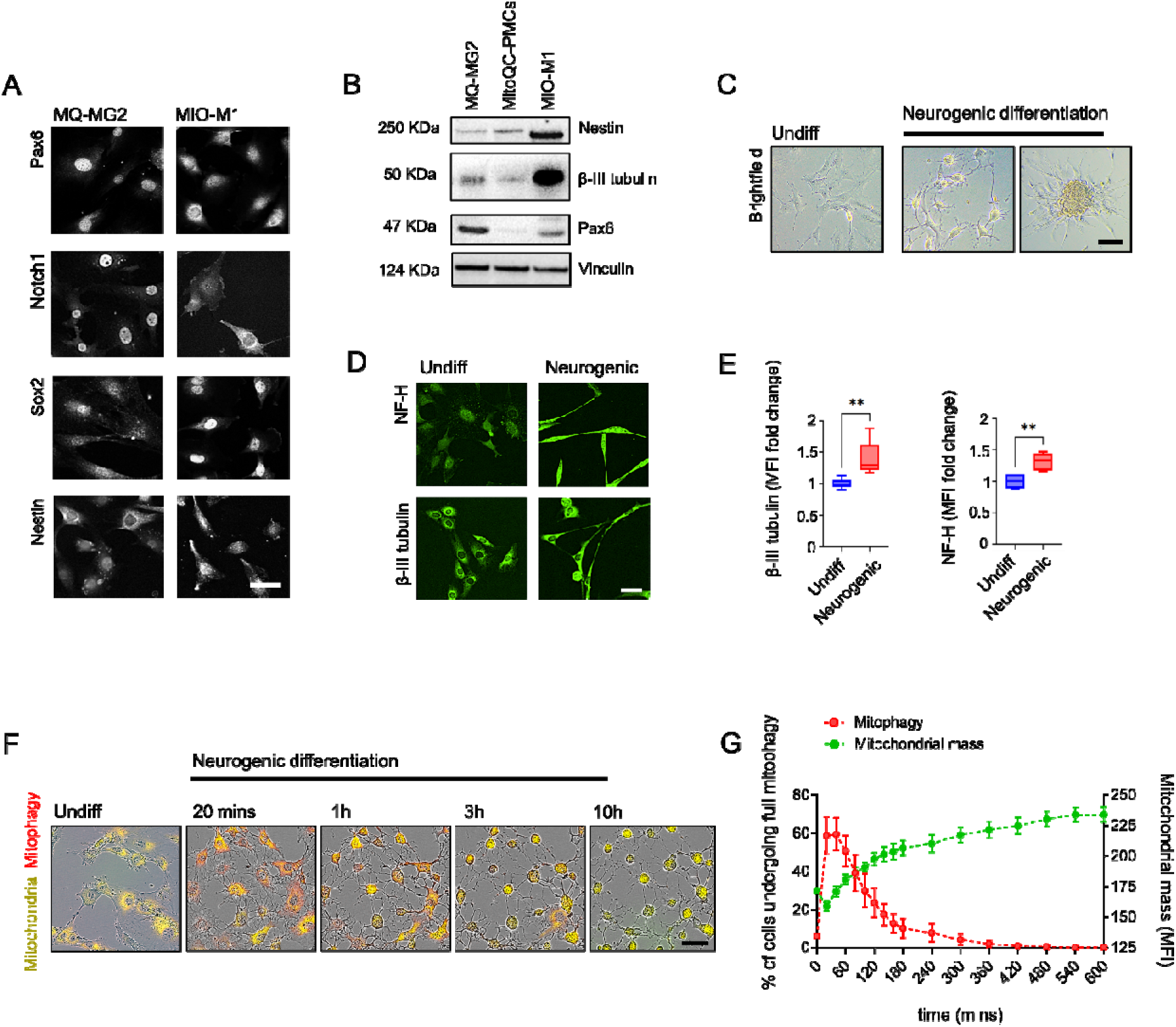
Neurogenic differentiation in MQ□MG2 Muller glia is supported by transient mitophagy and sustained mitochondrial biogenesis. **(A)** Representative micrographs of mouse MQ-MG2 and human MIO-M1 cells stained for canonical glial stem-cell markers (Pax6, Notch1, Sox2, and Nestin). **(B)** Representative immunoblots showing basal expression of glial stem-cell markers in lysates from MQ-MG2 and MIO-M1 Muller glial cell lines. Vinculin was used as the loading control. **(C)** Brightfield micrographs of MQ-MG2 cells under basal conditions and following neurogenic differentiation, demonstrating the emergence of complex neurite-like networks (3lJh) and neurosphere formation (4 days). **(D-E)** Representative micrographs and quantification of neurofilament-H (NF-H) and β-III tubulin expression in MQ-MG2 cells under basal conditions and after 3 days of neurogenic differentiation. **(F-G)** Representative IncuCyte® live-cell imaging and real-time kinetic analysis of mitophagy and mitochondrial mass over a 10-hour period (20-minute intervals) in MQ-MG2 cells undergoing neurogenic induction. Data are presented as box-and-whisker plots (median, interquartile range, and minimum/maximum values) (E); mean□±□SD (G). *n*□≥ 6 (E), *n*□≥ 4 (G). **P < 0.01. Unpaired Student’s *t*-test. Scale bars: 20 µm.

Previous studies have shown that mitophagy is critical for neuroblast differentiation into mature retinal ganglion cells, by facilitating the metabolic transition toward glycolysis prior the establishment of oxidative phosphorylation capacity [8]. However, whether Muller glial progenitors engage MQC to support neurogenic transitions remains unclear. Taking advantage of differentiation potential of MQ-MG2, *Incucyte* live-cell imaging revealed a transient, rapid upregulation of mitophagy within 30 minutes of neuronal induction (Figure 6F-G). This was followed by a sustained increase in mitochondrial mass, consistent with mitochondrial biogenesis (Figure 6F-G), which was confirmed by elevated TOMM20 and TFAM expression 48 hours post-differentiation (Figure 7A-B).

**Figure 7.**
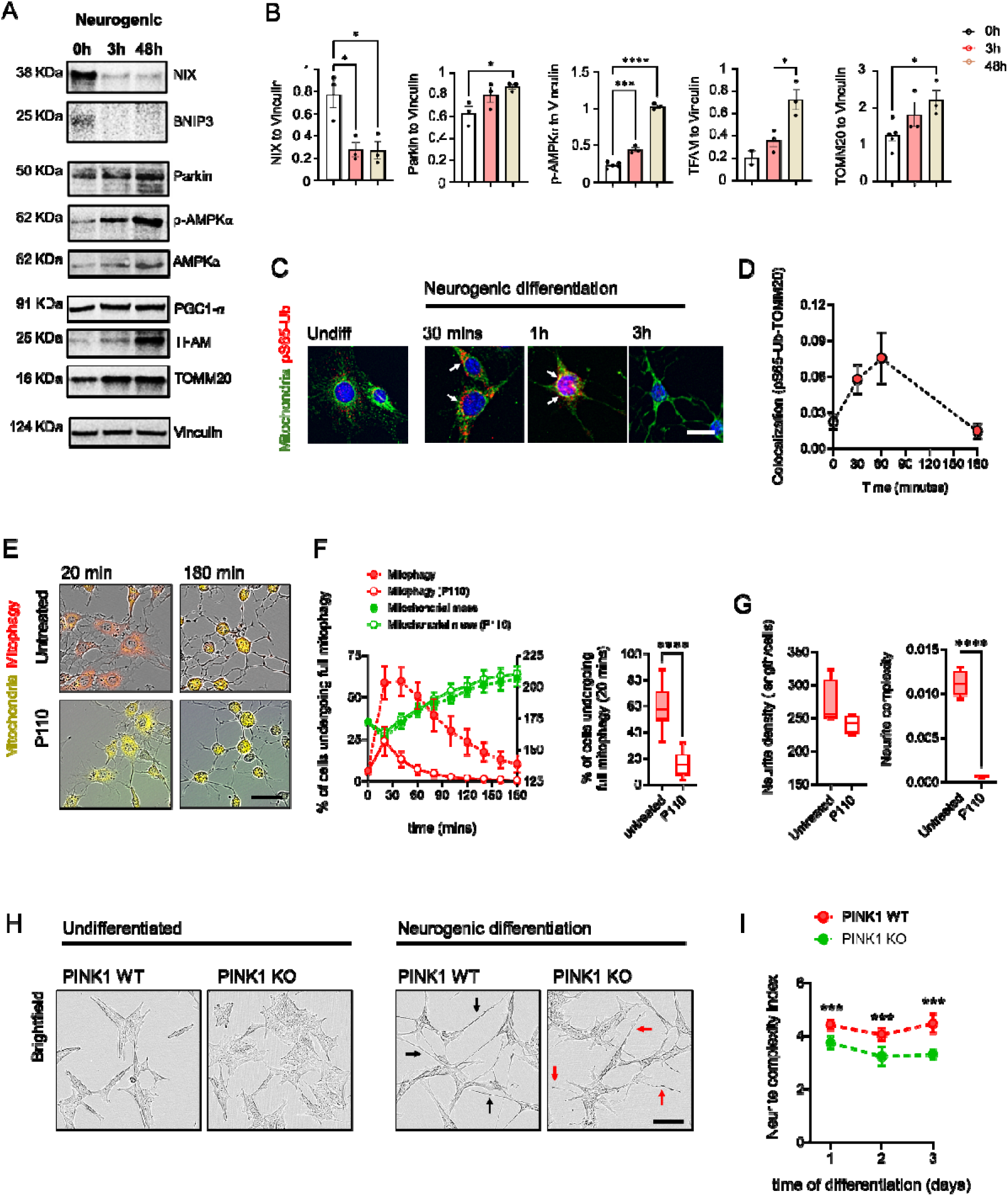
Fission-dependent and PINK1-linked mitochondrial remodelling supports neurite maturation in mammalian neurogenic systems. (A-B) Representative immunoblots and quantification of NIX, Parkin, phosphorylated AMPK (pThr172), TFAM, and TOMM20 at defined time points following neurogenic induction in MQ-MG2 cells. **(C-D)** Representative images and colocalization analysis of TOMM20 and phosphorylated ubiquitin (pS65-Ub), quantified using Pearson’s correlation coefficient. **(E-G)** MQ-MG2 cells were pre-treated with the mitochondrial fission inhibitor P110 peptide for 3 days prior to neurogenic induction. **(E-F)** IncuCyte® live-cell imaging and real-time kinetic analysis of mitophagy and mitochondrial mass over a 3-hour period (20-minute intervals) across treatment groups. The graph in (F) shows mitophagy levels at the peak of induction. **(G)** Quantification of neurogenic differentiation, assessed by neurite density and complexity (see “Methods” for calculation details) after 10□hours of neurogenic induction across treatment groups. **(H)** Brightfield micrographs of wild-type (WT) and PINK1 knockout (KO) SH-SY5Y cells under basal conditions and following 48 hours of neurogenic induction. PINK1-deficient cultures exhibit reduced neurite density and complexity (red arrows) compared with WT cells (black arrows). **(I)** Quantification of neurogenic differentiation, assessed by neurite density and complexity of neurogenic induction in WT and PINK1 KO cultures. Data are presented as mean□±□SD with individual data points (B); mean□±□SD (D, F, I); or box-and-whisker plots (median, interquartile range, minimum/maximum) (F-G). Protein levels were normalized to Vinculin. *n*□≥□3 (B-G); *n*□=□9 (I). *P < 0.05, ***P < 0.001, ****P < 0.0001. One-way ANOVA with Dunnett’s multiple comparison (B); unpaired Student’s *t*-test (F-G); Two-way ANOVA (I). Scale bars: 10 µm (E), 20 µm (A, G); 30 µm (I).

Immunoblotting and immunocytochemical analyses further revealed a rapid, biphasic engagement of mitophagy pathways during early neurogenic differentiation. Within 20–60 minutes of induction, both PINK1-dependent and receptor-mediated programmes were activated, as evidenced by peaks in mitochondrial pS65-Ub (Figure 7C-D) and BNIP3/NIX (Supplementary Figure 6), respectively. This early mitophagic phase transitioned into sustained AMPK activation (p-AMPK^Thr172^) and elevated Parkin expression by ∼3h (Figure 7A-B). In contrast, BNIP3 and NIX levels declined significantly from 1h onward (Supplementary Figure 6 and Figure 7A–B), consistent with transient engagement of receptor-mediated mitophagy during early stages, yet supressed as mitochondrial biogenesis becomes established. A similar temporal pattern was observed in human MIO-M1 cells (Supplementary Figure 7), supporting conserved metabolic remodelling whereby PINK1-dependent and receptor-mediated mitophagy are sequentially coordinated during Muller glial neurogenic transition.

To directly evaluate the contribution of mitophagy to glial-to-neuronal differentiation, MQ-MG2 were pre-treated with P110 (1 µM) for 3 days prior to neuronal induction. Live-cell imaging demonstrated that P110 markedly suppressed the early mitophagy response without preventing the subsequent increase in mitochondrial mass (Figure 7E-F). Although P110-treated cells still extended initial neurite-like projections, quantitative morphometric analysis revealed a significant reduction in neurite complexity (Figure 7E, G), indicating impaired neuronal maturation.

To obtain independent genetic evidence for PINK1 involvement, we differentiated a CRISPR-engineered PINK1 knockout SH-SY5Y cell line under comparable conditions. In contrast to wild-type controls, PINK1-deficient cells exhibited significantly reduced neurite complexity during differentiation, despite preserved viability under basal conditions (Figure 7H, I). Together, these findings indicate that fission-dependent mitophagy and PINK1-linked mitochondrial remodelling contribute to neurite maturation in mammalian neurogenic systems. Moreover, the data support a model in which distinct mitophagy pathways are temporally coordinated during Muller glial lineage transitions.

## DISCUSSION

Understanding how MQC is coordinated in the mammalian retina is essential for defining pathway-specific mechanisms of mitophagy dysregulation in disease. Advances in next-generation reporters, including mito-QC, have enabled spatially resolved interrogation of mitochondrial turnover *in vivo*. Using the MQ-MG2 model, we show that Muller glia function as important cellular nodes of MQC by engaging distinct mitophagy pathways under basal, metabolic stress, and differentiation conditions. Both PINK1-dependent and receptor-mediated mitophagy programs coexist within these cells and are differentially activated, positioning Muller glia not only as metabolic support cells but as active integrators of mitochondrial turnover capable of adapting mitophagy responses to physiological demand.

We previously demonstrated that mitophagy occurs in Muller glia within the outer retina [3]. Here, we extend these findings by showing that Muller glia harbor both PINK1-dependent and receptor-mediated (BNIP3/NIX) mitophagy pathways, comparable to those operating in other neuronal populations in the mouse and human retina. Importantly, these programs are retained in MQ-MG2 cells, which respond robustly to pathway-specific mitophagy inducers, a finding further supported by *in vivo* validation. Together, these observations reinforce Muller glia as important regulators of mitophagy within the retina, complementing evidence that they also participate in trans-mitophagy, a process described in zebrafish retina and analogous to mitochondrial transfer to astrocytes in the optic nerve [14, 32]. Collectively, these data support a model in which Muller glia coordinate both intrinsic and extrinsic MQC mechanisms to maintain retinal homeostasis.

Our pharmacological analyses demonstrate that Muller glia are capable of engaging both PINK1-dependent and PINK1-independent mitophagy pathways. Accordingly, KR, a potent activator of the PINK1–Parkin axis, and DFP, a well-established inducer of BNIP3/NIX-mediated receptor mitophagy, each elicited robust mitophagy responses *in vivo*. Consistent with scRNA-seq datasets revealing broad expression of receptor-mediated mitophagy machinery across outer retinal cell types, these findings suggest that PINK1-independent pathways may represent a prominent route of basal mitophagy in the retina [23]. This is further supported by the heightened sensitivity of mitophagy activation to DFP across multiple outer retinal layers. In contrast, KR produced a more spatially restricted response, prominently enhancing mitophagy at the OLM. Together, these observations indicate that distinct mitophagy programs are spatially segregated within the vertebrate retina, supporting a compartmentalized organization of MQC. Moreover, the coexistence of multiple mitophagy pathways in Muller glia may provide functional flexibility, enabling not only the targeted removal of damaged mitochondria via PINK1-mediated mechanisms, but also adaptive control of mitochondrial mass through receptor-mediated pathways under distinct physiological states such as hypoxia [33, 34].

The presence of multiple pharmacologically accessible mitophagy nodes highlights the mechanistic diversity through which mitochondrial homeostasis may be regulated under disease-relevant stress. This includes KR, which rescued mitochondrial dysfunction in MQ-MG2 cells under diabetic retinopathy–associated hyperfusion stress, consistent with our previous findings [3]. In addition, we identified mitophagy-enhancing agents acting in Muller glia through receptor-mediated pathways, including NIX-associated signalling activated by BDNF mimetics such as 7,8-DHF, as well as TSPO ligands (e.g., AC-5216). While activation of the BDNF/TrkB axis has previously been shown to induce NIX-dependent mitophagy [26], the molecular basis of TSPO-mediated mitophagy remains unclear. Notably, elevated TSPO signalling has been reported to suppress mitochondrial clearance downstream of the PINK1–Parkin pathway [35], whereas more recent studies suggest that TSPO ligands, including AC-5216, can attenuate mitophagy deficits in cellular models of Alzheimer’s disease [27]. Collectively, these findings reveal multiple pharmacologically accessible mitophagy routes in Muller glia and provide a framework for future investigation of mitophagy regulation under pathological stress.

One of the central findings of this study is the identification of fission-dependent mitophagy as a regulatory node during the early stages of Muller glial differentiation. Using MQ-MG2 cells, we observed a rapid mitophagy burst within 30 minutes of neurogenic induction, followed by coordinated mitochondrial biogenesis, consistent with a transition from mitochondrial clearance to establishment of oxidative phosphorylation capacity. Pharmacological inhibition of fission-dependent mitophagy using P110 suppressed the early mitophagy pulse, disrupted mitochondrial remodelling, and reduced neurite complexity, indicating that fission-linked mitochondrial turnover contributes to neurogenic maturation. Previous studies have linked mitophagy to retinal ganglion cell maturation, primarily through NIX-dependent pathways that promote a glycolytic shift [8, 36, 37]. More recently, both PINK1- and NIX-mediated mitophagy have been implicated in embryonic retinal development [38].

Our findings extend this framework by demonstrating that distinct mitophagy pathways are temporally coordinated during Muller glial lineage transitions. PINK1-dependent activation remained sustained throughout neurogenic progression, whereas NIX-mediated mitophagy followed a transient, biphasic pattern that was later attenuated. These observations support a model in which NIX-associated mitophagy contributes to early mitochondrial clearance and is later attenuated as mitochondrial biogenesis proceeds, while PINK1-dependent mitophagy may support sustained MQC during differentiation. This interpretation is further supported by our pharmacological evidence, where NIX activation via DFP or 7,8-DHF reduced mitochondrial mass, while PINK1 activation with KR induced mitophagy without depletion of mitochondrial content, suggesting compensatory coupling to mitochondrial biogenesis. The functional relevance of PINK1□dependent mitophagy for neurogenic differentiation was further supported by reduced neurite complexity in PINK1□deficient SH□SY5Y cells, indicating that PINK1 may contribute to neuronal maturation across distinct neurogenic contexts. Together, these findings suggest that coordinated mitophagy pathways regulate adult glial-to-neuronal transitions by shaping mitochondrial turnover during cellular remodelling.

Beyond the mechanistic insights described here, MQ-MG2 provides a tractable experimental system for interrogating mitochondrial quality control in Muller glia. Unlike transient mitophagy reporter systems or primary Muller cultures, MQ-MG2 combines stable mito-QC expression with preservation of endogenous mitophagy adaptors, neurogenic competence, and metabolic features relevant to retinal physiology. This enables pathway- and time-resolved analysis of mitochondrial turnover without genetic overexpression, facilitating investigation of MQC under both physiological and pathological conditions. In addition, MQ-MG2 overcomes limitations associated with primary cultures, including senescence, variability, and restricted scalability. The compatibility of Mito-QC with fixation and multiplexed imaging [15] further supports integrative analysis of mitochondrial dynamics while reducing reliance on *in vivo* experimentation.

Collectively, our findings support Muller glia as active regulators of mitophagy and demonstrate that PINK1-dependent and receptor-mediated pathways can be selectively activated in a context-dependent manner. We further implicate PINK1-linked mitochondrial remodelling as an early regulatory component of Muller glial neurogenic transitions. These mechanistic insights advance our understanding of how pathway-specific mitophagy may contribute to retinal mitochondrial homeostasis, regenerative potential, and disease-associated mitochondrial dysfunction.

## MATERIALS AND METHODS

### Animals

Male and female Mito-QC^+/+^ mice on a C57BL/6J background were used in this study. The Mito-QC reporter incorporates a tandem mCherry–GFP fluorescent tag targeted to mitochondria via a signal peptide derived from the outer mitochondrial membrane protein Fis1 [15]. Detection of the Mito-QC-knockin allele (mCherry-GFP-mtFIS1^101–153^) was determined by PCR [15]. Mice were maintained under standard specific pathogen–free conditions at 20–24□°C and 45–65% relative humidity, with a 12□h light/dark cycle and *ad libitum* access to food and water. All animal experiments were approved by the Ethics Committees of the University of Birmingham (Project Licence PP6860623) and Queen’s University Belfast (Project Licence PPL2814). Procedures were reviewed by the local Animal Welfare and Ethical Review Bodies (AWERB) and conducted in accordance with the UK Animals (Scientific Procedures) Act 1986. Animal use conformed to the Association for Research in Vision and Ophthalmology (ARVO) Statement for the Use of Animals in Ophthalmic and Vision Research.

### Animal treatments

Fourteen-week-old Mito-QC^+/+^ mice were treated for one week by supplementation of Kinetin Riboside (60□mg/L), deferiprone (1mg/L and 10 mg/L), 7,8-dihydroxyflavone (20 mg/L) or vehicle control (0.1% DMSO) in the drinking water. Treatments were administered three times per week on alternating days (Days 1, 3, and 5) of the treatment phase. Age-matched Mito-QC^+/+^ mice receiving no treatment were included as untreated controls.

### Retinal immunohistochemistry (IHC)

Mice were euthanised by CO□ inhalation, and eyes were rapidly enucleated and fixed in 4% paraformaldehyde (Sigma-Aldrich, Dorset, UK) for 2□h at room temperature. Eyes were then processed for IHC as previously described [39, 40]. Briefly, 14 µm-thick retinal cryosections were blocked in 10% normal donkey serum (NDS) in phosphate-buffered saline (PBS) and incubated overnight at 4□°C with rabbit anti–glutamine synthase antibody (1:10,000; G2781, Sigma-Aldrich) diluted in PBS containing 0.5% Triton X-100 and 10% NDS. Following incubation, sections were washed in PBS and incubated for 1□h at room temperature with donkey anti-rabbit Alexa Fluor™ Plus 405 secondary antibody (1:400; Thermo Fisher). Sections were then mounted using Vectashield (Vector Labs, Peterborough, UK) and imaged by confocal microscopy (C1 Nikon Eclipse TE200-U, Nikon UK Ltd, Surrey, UK). For quantification of mitophagy, retinal sections were mounted without immunohistochemical staining.

### Isolation of Mito-QC Primary Muller Cells

Mito-QC Primary Muller Cells from 8-week old male and female Mito-QC^+/+^ mice were isolated following previously established methods [2, 41]. In brief, retinas were dissected and rinsed in Hanks’ Balanced Salt Solution (HBSS) before incubation with 2% dispase in a humidified 5% CO□ incubator at 37□°C for 1□h. Enzymatic activity was neutralised by washing with Dulbecco’s Modified Eagle Medium (DMEM) containing 5.5□mM D-glucose and 10% fetal bovine serum (FBS). Retinas were then transferred to 6-well culture plates and maintained at 37□°C in 5% CO□ until confluence was reached. After two to three passages, PMC cultures achieved high relative purity, as assessed by characteristic cellular morphology and confirmed by immunostaining for Muller glia markers, including glutamine synthetase.

### Monoclonal Isolation of Spontaneously Immortalised Mito-QC Cell Lines

A heterogeneous population of spontaneously immortalised cells capable of proliferating beyond 30 passages was derived from cultured mito-QC PMCs. Monoclonal cell lines were generated by single-cell cloning using serial dilution after passage 30, according to established protocols [42, 43]. To support cell survival and proliferation, sterile-filtered conditioned medium was added to each well. Wells containing single-cell–derived colonies were identified and marked, and cultures were maintained for 18 days at 37□°C in a humidified 5% CO□ incubator. Eight independent clones were subsequently selected and expanded for further characterisation.

### Cell culture

Mito-QC PMCs, immortalized Mito-QC Muller glia clones, the human Muller cell line MIO-M1 (obtained from the UCL Institute of Ophthalmology), and SH-SY5Y neuroblastoma cells (wild type and CRISPR–PINK1 knockout; AB280876, Abcam) were used in this study. Murine and human Muller glial cell lines were cultured in DMEM containing normal glucose (5.5□mM), supplemented with 10% heat-inactivated FBS and 1% penicillin/streptomycin. Cells were maintained at 37□°C in a humidified incubator with 5% CO□. SH-SY5Y cells were cultured according to the manufacturer’s instructions in a 1:1 mixture of Eagle’s Minimum Essential Medium (EMEM) and Ham’s F-12K medium, supplemented with 10% FBS and 1% penicillin/streptomycin under normal-glucose conditions.

Cells were routinely maintained in T75 flasks and passaged every 3–4 days upon reaching 70–80% confluence. Mito-QC PMCs were used for experiments between passages 2 and 4. For *in vitro* modeling of diabetic conditions, cells were cultured in DMEM supplemented with 30.5□mM D-glucose (high glucose; HG) for 3 days, as previously described [3]. No mycoplasma contamination was detected.

### Cell Growth Characteristics

The growth rate of MQ-MG2 cells was assessed using previous protocols [43]. Briefly, cells were seeded at a density of 10,000 cells per well in 24-well plates and maintained under physiological glucose conditions (5.5□mM). Culture medium was replaced every two days. Cell counts were performed daily for seven consecutive days using a haemocytometer, with measurements taken from three independent wells per time point. The mean cell number was used to calculate the mean growth rate (MGR).

### Cell Treatments

Cells were seeded into 96-well plates and allowed to adhere overnight before treatment with compounds of interest for 16□h. Mitophagy-modulating agents included kinetin (0.3–5□µM; Sigma-Aldrich), Kinetin Riboside (0.3–5□µM; Sigma-Aldrich), niclosamide (1□µM; Bio-Techne), urolithin-A (50□µM; Bio-Techne), 7,8-dihydroxyflavone (0.1–50□µM; Cayman Chemical), AC-5216 (0.01–50□µM; Cayman Chemical), and Deferiprone (1–1000□µM). Where relevant, compounds were prepared in the manufacturer-recommended solvents, including 0.1% DMSO as vehicle control. For induction of mitophagy by nutrient deprivation or hypoxia, cultures were maintained in HBSS or under low-oxygen conditions (1% O□) for 24 hours.

To antagonise mitochondrial fission, cells were pre-incubated with the DRP1-inhibitory peptide P110 (1□µM; Bio-Techne) [44] for 2□h under normal-glucose. Cultures were then washed in PBS and maintained in HG supplemented with P110 (1□µM) for 72□h. Additional treatments with Kinetin Riboside (100 nM) or vehicle (0.1% DMSO) were applied under HG ± P110 conditions. Treatments were changed daily and lasted 72□h before final experimental readouts.

### Immunocytochemistry

Cells were fixed in 4% PFA, rinsed in PBS, permeabilized with 0.5% Triton X-100, and blocked with 10% FBS in PBS. Cells were then incubated overnight at 4□°C with primary antibodies (Table S2) diluted in PBS containing 10% FBS and 0.05% Tween-20 (Sigma-Aldrich). After incubation, cells were rinsed in PBS, probed with appropriate fluorophore-conjugated secondary antibodies for 90 min at room temperature, washed again in PBS, and counterstained with DAPI nuclear dye (1:1000; Sigma-Aldrich) prior to microscopy imaging.

### Western Blotting

Cells were washed with ice-cold PBS, scraped into ice-cold RIPA buffer, and incubated on ice for 30 min. Lysates were centrifuged at 13,500 × g for 5 min at 4 °C, and supernatants were passed through a QIAshredder column to remove genomic DNA. Protein concentration was determined by Bradford assay. Samples were diluted in deionised water, mixed with XT sample buffer and reducing agent, and heated at 95 °C for 5 min. Equal amounts of protein (20–50 µg) were resolved on XT Bis-Tris gels using XT MES running buffer at 100 V for 90 min and transferred to PVDF membranes using the Bio-Rad Trans-Blot Turbo system. For PINK1 immunoblotting, proteins were separated on NuPAGE Bis-Tris gels with MOPS running buffer and transferred to PVDF membranes using a wet-transfer system. Membranes were blocked with 5% milk in TBST for 1 h and incubated overnight at 4 °C with primary antibodies (Table S2) diluted in 3% BSA/TBST. After washing, membranes were incubated with HRP-conjugated secondary antibodies for 90 min at room temperature. Signals were visualised using Immobilon chemiluminescent substrate and imaged on a ChemiDoc™ imaging system (Bio-Rad). Band intensities were quantified by densitometry in FIJI (ImageJ) software (NIH, Bethesda, MD, USA).

### Extracellular metabolic flux analysis

OCR and PER were obtained using the Seahorse XFe-96 Extracellulr Flux Analyzer (Agilent Technologies, Stockport, UK). Cells were seeded in Seahorse XFe96 microplates at 1,200 cells/well (MIO-M1) or 4,000 cells/well (MQ-MG2 and Mito-QC PMCs), allowed to adhere at room temperature, and incubated at 37□°C. Treatments were refreshed daily and maintained for 72□h prior to metabolic assessment. Cells were washed and medium replaced with fresh phenol red–free XF DMEM with glucose (7□mM), sodium pyruvate (1□mM), HEPES (5□mM) and L-glutamine (2□mM). A standard *Mito Stress Test* was performed with sequential injections of oligomycin (2 µM), FCCP (2 µM), and rotenone plus antimycin-A (0.5 µM each), followed by Hoechst (1 µg/mL). A modified *Mito Stress Test* was also conducted to obtain glycolytic rate measurements. Injections comprised oligomycin (2 µM), BAM15 (3 µM), rotenone plus antimycin-A (2 µM each), and monensin (25 µM) with Hoechst (1 µg/mL). OCR and PER were analysed using Seahorse Wave software (Agilent), with well normalisation based on Hoechst mean fluorescence intensity (MFI) measured on a BMG CLARIOstar plate reader.

### Neuronal Differentiation

Muller glial (MQ-MC2 and MIO-M1) and SH-SY5Y cells were induced toward neuronal differentiation using adapted protocols [31]. Cells were seeded onto fibronectin-coated culture plates (10□ng/mL, 1□h at room temperature) and maintained for 24□h prior to induction. Differentiation was initiated by replacing growth medium with commercial neurogenic differentiation medium (PromoCell) for a maximum of 3 days. Neuronal differentiation was confirmed by immunostaining for mature neuronal markers, including neurofilament-H and β-III tubulin, together with the formation of neurite networks.

### Imaging Morphometry

Epifluorescence images were acquired using an Olympus IX7 inverted microscope with Image-Pro Plus software, using identical acquisition settings across experimental conditions. Time-lapse imaging was performed using the *IncuCyte*® Live-Cell Analysis System (Sartorius, Surrey, UK), with images acquired automatically every 20–30□min for up to 3 days. For retinal analyses, confocal images were acquired using a Nikon C1 confocal microscope (Eclipse TE200-U; Nikon) and Nikon EZ-C1 software, with photomultiplier tube settings kept constant across samples. Image analysis was performed using FIJI (ImageJ). For cell culture experiments, epifluorescence images were selected from equivalent cardinal regions of each well, guided by bright-field or DAPI nuclear signals. Background subtraction was applied uniformly to all images prior to quantitative analysis.

#### Assessment of Mitophagy (Mito-QC) and mitochondrial biogenesis in fixed samples

Quantification of mitophagy in MQ-MG2 cells, Mito-QC PMCs, and Mito-QC retinal sections was performed as previously described [2, 3]. Briefly, GFP signal was subtracted from the mCherry signal within defined regions of interest using the *Image Calculator* function in FIJI (ImageJ). The resulting image, representing mitolysosomes, was thresholded and binarized using identical parameters across all experimental groups, and total mitolysosome area was quantified by particle analysis. Mitophagy measurements were normalized to mitochondrial mass, estimated from the FIS1–GFP signal. For TFAM and Cox4 analyses, mean fluorescence intensity (MFI) was quantified within the OLM–OPL region, and background fluorescence was subtracted from non-stained retinal areas. For cell-based analyses, quantification was performed on more than 50 cells per independent biological replicate.

#### Assessment of Mitophagy and mitochondrial biogenesis using IncuCyte® Live-Cell

Mitophagy in MQ-MG2 cells was quantified by a blinded assessor by scoring the proportion of cells undergoing partial or complete mitophagy, normalized to total cell density per field of view. For assessment of mitochondrial biogenesis, GFP fluorescence was quantified over time by calculating the MFI per cell at each time point following background subtraction.

#### Co-localization analysis

Following immunostaining for TOMM20 in combination with pS65-Ub, cells were analysed using the Colocalization Threshold module in FIJI to obtain Pearson’s correlation coefficient.

#### Ki67 (nuclear) and neurogenic marker analysis

For Ki67 analysis, DAPI-positive signals were binarized using constant threshold values across all experimental groups and processed for particle analysis. Resulting region of interests (ROIs) were then overlaid onto Ki67 to obtain MFI per nuclei following background subtraction. For analysis of neurofilament-H and β-III tubulin, individual cells were manually traced, and MFI was quantified after background subtraction.

#### Neurite quantification

Bright-field images were skeletonized and analysed to quantify (i) total neurite length, (ii) average neurite length, (iii) neurite junction density, and (iv) neurite endpoint density. Neurite density was calculated by normalizing total neurite length to cell number per field of view. Neurite complexity was expressed as a reticulated structural index, calculated using the following formula:

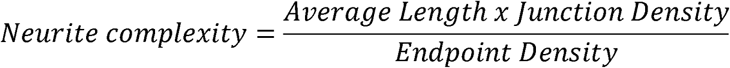

### Statistical Analysis

Comparisons between two experimental groups were performed using unpaired Student’s *t*-tests. For analyses involving more than two groups, one-way or two-way analysis of variance (ANOVA) followed by *post hoc* test was applied (as detailed in figure legends). Statistical outliers were identified and removed using Grubb’s test (α = 0.05). For retinal analyses, 6–8 images (derived from at least two retinal sections per eye) were quantified, and values were averaged per eye before statistical comparison. The animal sample size was adjusted to *n*□=□3–7 eyes in each mouse group. Single-cell RNA-seq transcriptomic data from previously published, pre-processed, and annotated datasets were retrieved from the CZ CELLxGENE project to assess expression of mitophagy adaptor genes in adult human (Wang *et al.*, 2022) [16] and mouse retina (Li *et al.*, 2024) [17]. No additional normalization or reprocessing of raw sequencing data was performed. Principal component analysis (PCA) with confidence ellipses for bioenergetic parameters was performed using SRplot (https://www.bioinformatics.com.cn/en). Data are presented as mean□±□SD or mean□±□SE or as box-and-whisker plots, where the median is shown as the central line, box boundaries represent the interquartile range (IQR), and whiskers represent the minimum and maximum values within 1.5× the IQR. A *P*-value <□0.05 was considered statistically significant. All statistical analyses were performed using GraphPad Prism (v10.1.0, La Jolla, CA).

## Acknowledgments

Special thanks to Ian Ganley (University of Dundee) for providing us with the *mitoQC* mice and to Astrid Limb (University College London) for providing the MIO-M1 cell line.

## Data availability

The data that support the findings of this study are available from the corresponding author, JRH, upon reasonable request.

## Abbreviations

7,8-DHF: 7,8-dihydroxyflavone
BDNF: brain derived neurotrophic factor
BNIP3: BCL2/adenovirus E1B 19 kDa interacting protein 3
DFP: Deferiprone
Fis1: Mitochondrial Fission 1 protein
KR: Kinetin Riboside
MIO-M1: Moorfields/Institute of Ophthalmology-Muller 1
MQC: mitochondrial quality control
NIX: BCL2/adenovirus E1B 19-kDa-interacting protein 3-like
OCR: Oxygen consumption rate
PER: Proton efflux rate
PINK1: PTEN-induced kinase 1
PMC: Primary Muller cells
pS65-Ub: Phospho-S65-ubiquitin
TFAM: mitochondrial transcription factor A
TSPO: Translocator protein.

