## Supplemental information for "Pathway-selective mitophagy regulates retinal physiology and neurogenic transitions in Muller glia"

**SUPPLEMENTARY INFORMATION**

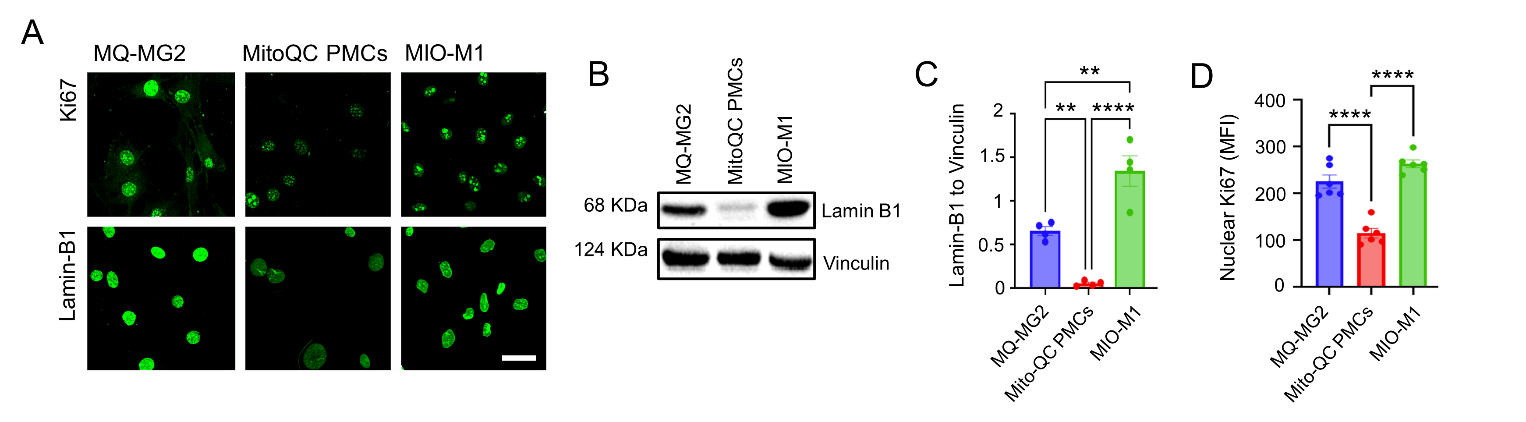

**Supplemental Fig 1. MQ-MG2 Müller glia bypass cellular senescence. (A)** Representative micrographs of MQ‑MG2, Primary Müller cells from Mito‑QC reporter mice (Mito‑QC PMCs) and human MIO‑M1 Müller cells stained for Ki67 and Lamin B1. **(B-C)** Immunoblot analysis and quantification of Lamin B1 in lysates from MQ‑MG2 cells, Mito‑QC PMCs, and human MIO‑M1 Müller cells. Protein levels were normalized to Vinculin. **(D)** Quantification of Ki67 mean fluorescence intensity (MFI) across Müller glial groups. Data are presented as mean ± SD. *n* ≥ 4. **P < 0.01, ****P < 0.0001. One-way ANOVA with Dunnett’s multiple comparison. Scale bar 20 µm.

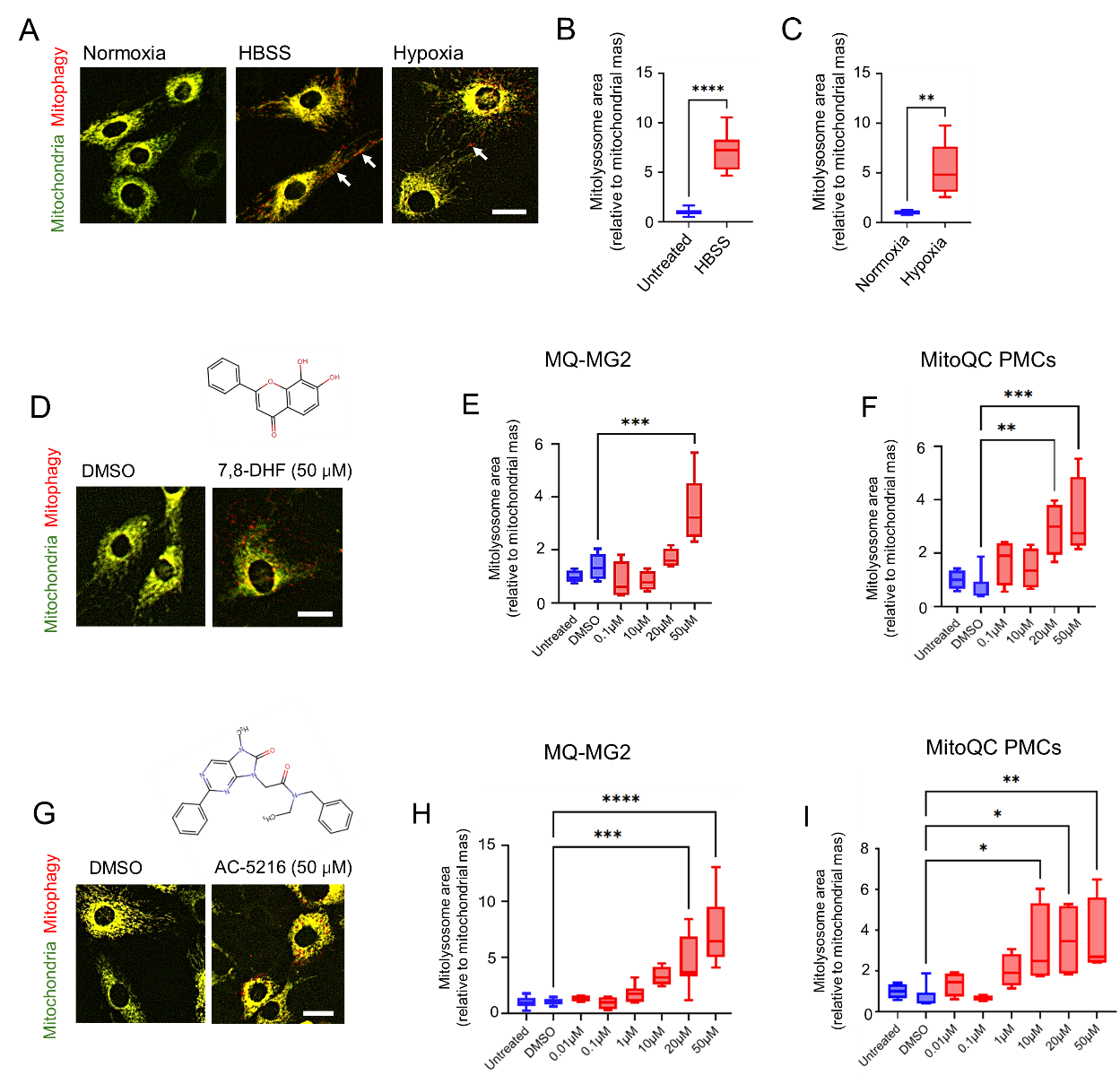

**Supplemental Fig 2. Mitophagy responsiveness of MQ-MG2 Müller glia to physiological stress and non-PINK1 pharmacological induction. (A-C)** Representative images and quantification of mitophagy (mCherry‑only puncta, arrows) in MQ‑MG2 following amino acid starvation (HBSS) or hypoxia (1% O₂) for 24h. **(D-I)** MQ‑MG2 were treated with different pharmacological agents, including **(D-F)** the BDNF mimetic 7,8-dihydroxyflavone (7,8-DHF) and **(G-I)** the TSPO ligand AC-5216. Responses were compared to primary Müller cells obtained from Mito-QC mice (Mito-QC PMCs). DMSO (0.1%) served as the vehicle control for all pharmacological treatments. Data are presented as box‑and‑whisker plots (median, interquartile range, and minimum/maximum values). *n* ≥ 4. *P < 0.05, **P < 0.01, ***P < 0.001, ****P < 0.0001. One-way ANOVA with Dunnett’s multiple comparison. Scale bars: 10 µm.

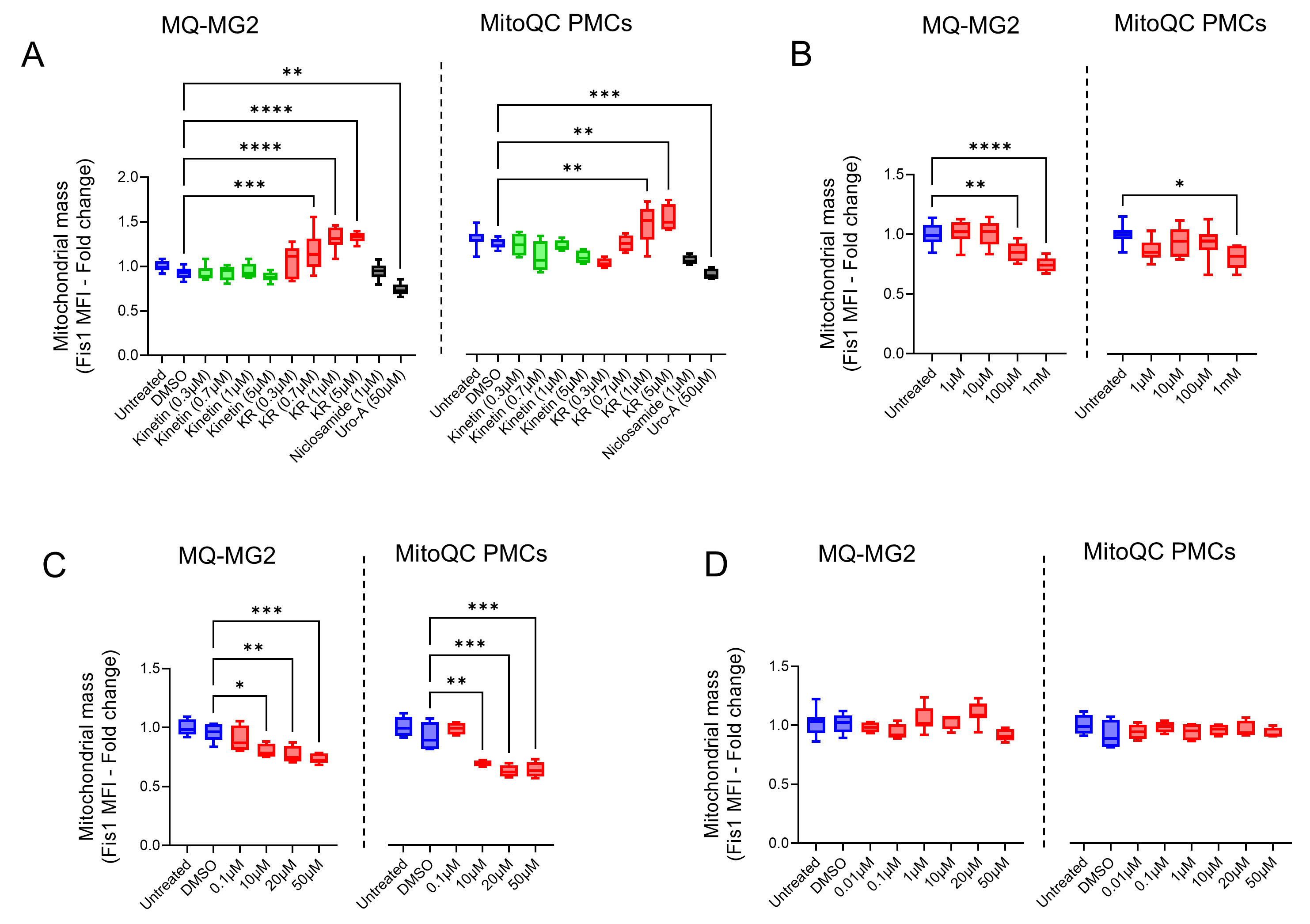

**Supplemental Fig 3. Mitochondrial mass profiles in MQ‑MG2 cells and Mito‑QC Primary Müller Cells (PMCs) following PINK1‑dependent and ‑independent mitophagy induction.** Quantification of mitochondrial mass (Fis1 mean fluorescence intensity [MFI]) in MQ‑MG2 cells and Mito‑QC PMCs after 24 h treatment with pathway‑selective mitophagy inducers, including **(A)** PINK1-dependent: Kinetin, Kinetin Riboside [KR], Niclosamide, and Urolithin A [Uro A]; **(B)** Deferiprone (iron chelator); **(C)** 7,8-dihydroxyflavone (BDNF mimetic); **(D)** AC-5216 (TSPO ligand). DMSO (0.1%) served as the vehicle control for all pharmacological treatments except Deferiprone. Data are presented as box‑and‑whisker plots (median, interquartile range, and minimum/maximum values). *n* ≥ 4. *P < 0.05, **P < 0.01, ***P < 0.001, ****P < 0.0001. One-way ANOVA with Dunnett’s multiple comparison.

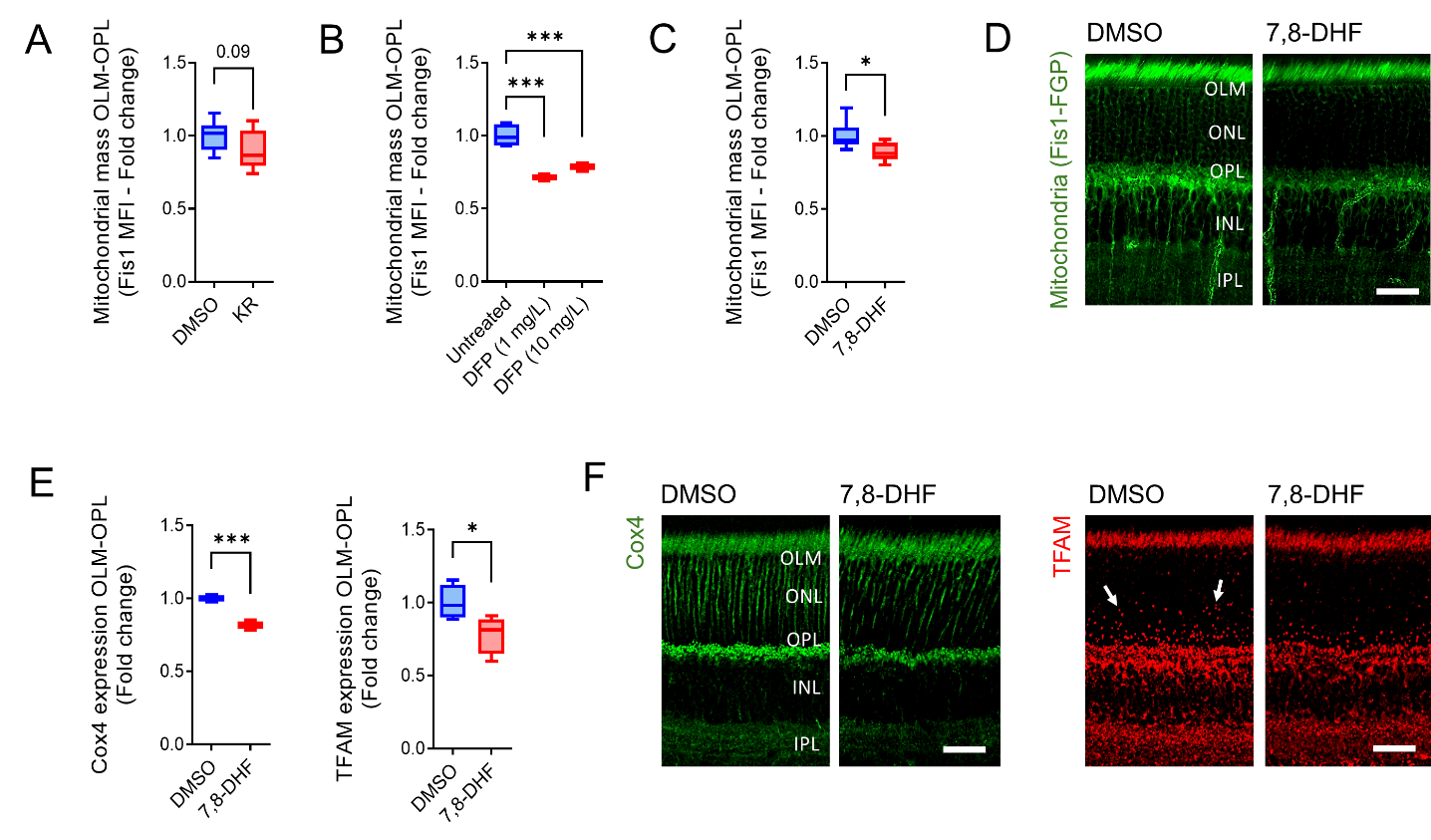

**Supplemental Fig 4. Differential modulation of mitochondrial mass by PINK1‑ and NIX‑dependent activators in the mouse retina.** Fourteen‑week‑old Mito-QC mice (A–D) or non‑Mito-QC mice (E, F) were treated for one week with Kinetin Riboside (KR; 60 mg/L), 7,8‑dihydroxyflavone (7,8‑DHF; 20 mg/L), Deferiprone (DFP; 1 mg/L or 10 mg/L), or vehicle control (0.1% DMSO) delivered in drinking water. **(A-D)** Quantification of mitochondrial mass in the outer retina based on Fis1‑GFP mean fluorescence intensity (MFI) across treatment groups. **(E-F)** Quantification of Cox4 and mitochondrial transcription factor‑A (TFAM) levels in the outer retina of 7,8‑DHF– and vehicle‑treated mice. Arrows in (F) indicate TFAM‑positive mitochondrial nucleoids. Data are presented as box‑and‑whisker plots (median, interquartile range, and minimum/maximum values). (A) *n* ≥ 7; (B) *n* ≥ 3; (C) *n* ≥ 6; (E) *n* = 4 eyes. *P < 0.05, ***P < 0.001. (A, C, E) Unpaired Student’s *t*‑test; (B) One-way ANOVA with Dunnett’s multiple comparison. OLM, outer limiting membrane; ONL, outer nuclear layer; OPL, outer plexiform layer; INL, inner nuclear layer; IPL, inner plexiform layer. Scale bars: 20 µm.

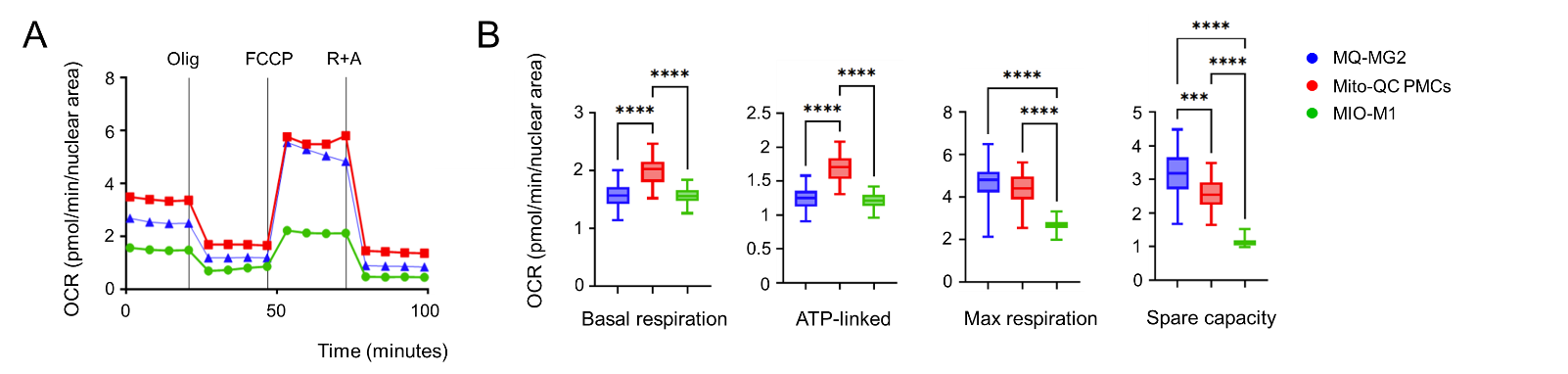

**Supplemental Fig 5. MQ-MG2 retain mitochondrial respiratory competence compared with primary Müller cells. (A)** Representative Seahorse extracellular flux traces from *Cell Mito Stress Test* performed on MQ‑MG2 cells, primary Müller cells isolated from Mito-QC reporter mice (Mito-QC PMCs), and human MIO‑M1 Müller cells. **(B)** Quantification of oxygen consumption rate (OCR) under basal and stressed conditions. Data are presented as box‑and‑whisker plots (median, interquartile range, and minimum/maximum values). *n* ≥ 14. ***P < 0.001, ****P < 0.0001. One-way ANOVA with Dunnett’s multiple comparison. Olig, oligomycin; R+A, rotenone + antimycin.

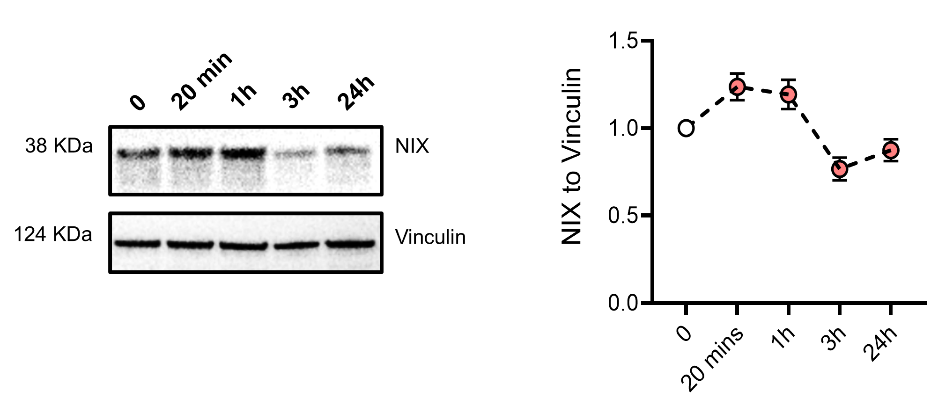

**Supplemental Fig 6. Transient upregulation of NIX in MQ‑MG2 Müller glia during neurogenic differentiation.** Representative immunoblots and quantification of NIX protein levels at defined time points following neurogenic induction in MQ‑MG2 cells. Protein expression was normalized to Vinculin. Data are presented as mean ± SE (*n* ≥ 4).

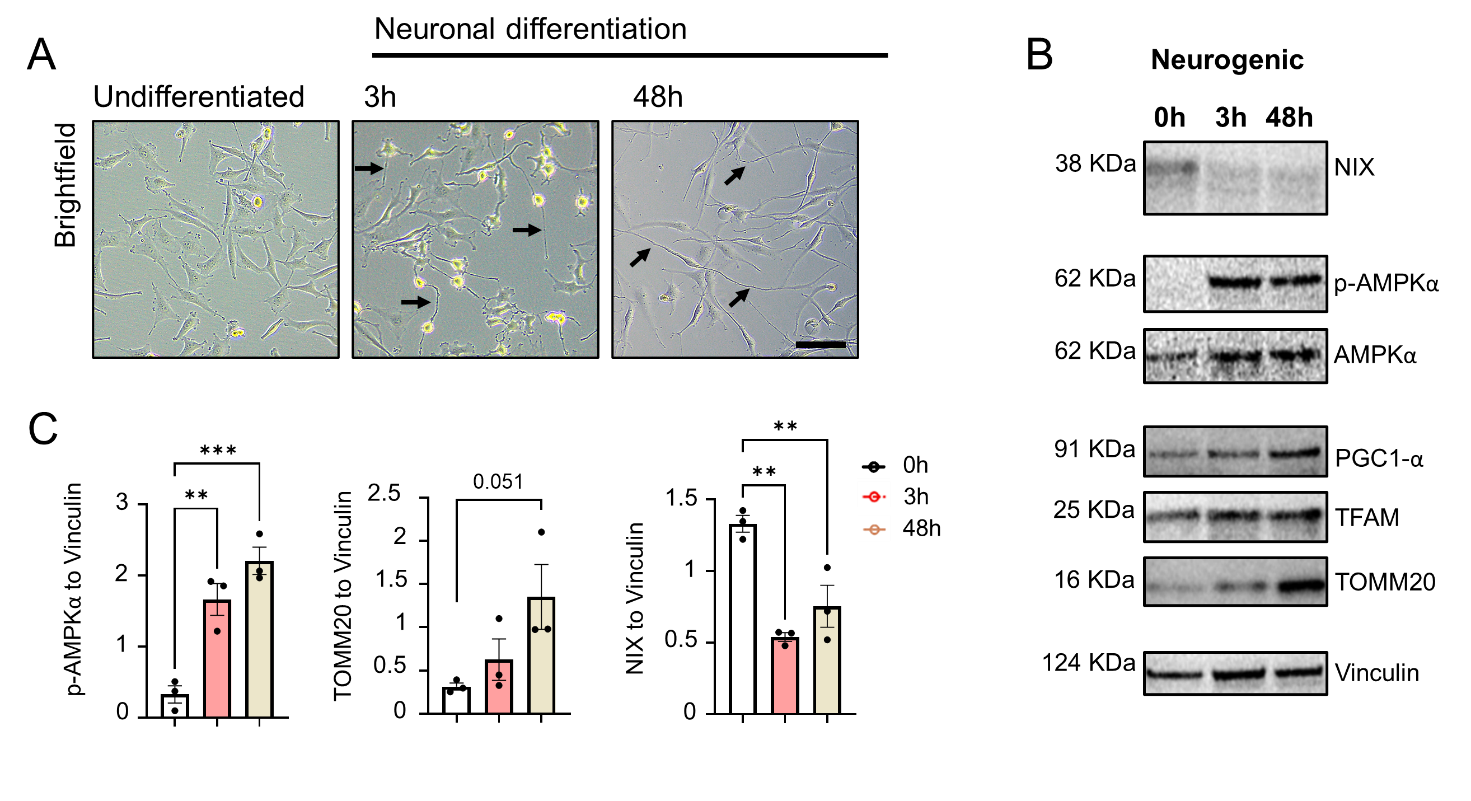

**Supplemental Fig 7. AMPK activation and NIX suppression during neurogenic differentiation of MIO‑M1 cells. (A)** Brightfield micrographs of MIO‑M1 cells under basal conditions and following neurogenic induction, showing the emergence of early neurite‑like extensions (3 h) and established neurite‑like networks (48 h). **(B-C)** Representative immunoblots and quantification of phosphorylated AMPK (pThr172), NIX and TOMM20 at defined time points following neurogenic induction. Data are presented as mean ± SD with individual data points. *n* ≥ 3. *P < 0.05, **P < 0.01, ***P < 0.001. One-way ANOVA with Dunnett’s multiple comparison. Scale bar 50 µm.

**TABLE S1. Phenotypic characterization of MQ‑MG2 relative to primary Müller cells (PMCs) and the human MIO‑M1 Müller cell line.**

| **Marker** | **Cell type** | **MQ-MGC2** | **PMCs** | **MIO-M1** |
| --- | --- | --- | --- | --- |
| GS | Müller | + | + | + |
| GFAP | Müller | + | + | + |
| α-SMA | Müller | + | + | + |
| S100β | Müller | + | + | + |
| AQP4 | Müller | + | + | + |
| Vimentin | Müller | + | + | + |
| IL-33 | Müller | + | + | NE |
| Kir4.1 | Müller | + | + | + |
| HO-1 | Müller | + | NE | + |
| S100α | Müller | - | - | - |
| CRALBP | Müller | - | - | + |
| Cone arrestin | Cone photoreceptors | + | + | + |
| Calbindin | Horizontal/amacrine | - | - | - |
| Iba-1 | Microglia | - | - | - |
| PCK-α | Rod Bipolar | + | + | + |
| Rhodopsin | Rod photoreceptors | + | + | NE |
| Nestin | Retinal stem cells | + | + | + |
| Pax6 | Retinal stem cells | + | + | + |
| Sox2 | Retinal stem cells | + | + | + |
| Notch1 | Retinal stem cells | + | + | + |
| β-III-tubulin | Mature neurons | + | + | + |
| Neurofilament-H | Mature neurons | + | NE | + |

(+) Expressed; (-) non-expressed; (NE) Not evaluated in this study

**TABLE S2. Primary antibodies used for western blot (WB) and immunocytochemistry (ICC).**

| **Antigen** | **Antiserum (Host)** | **Supplier** | **Cat. Number** | **Dilution** |
| --- | --- | --- | --- | --- |
| AMPK | rabbit | Cell Signaling | 2532S | 1:1000 (WB) |
| p-AMPKThr172 | rabbit | Cell Signaling | 2535S | 1:1000 (WB) |
| ATP synthase V | Mouse | Thermo Fisher | A-21351 | 1:1000 (WB) |
| AQP4 | Rabbit | Proteintech | 16473-1-AP | 1:1000 (WB) |
| α-SMA | Rabbit | Abcam | Ab32575 | 1:500 (ICC)  1:1000 (WB) |
| β-III-tubulin | Rabbit | Abcam | Ab18207 | 1:500 (ICC)  1:1000 (WB) |
| Β-Actin | Mouse | Sigma-Aldrich | A1978 | 1:10000 (WB) |
| BNIP3 | Mouse | Proteintech | 68091-1-Ig | 1:5000 (WB) |
| Calbindin | Rabbit | Swant | CB-38a | 1:500 (ICC) |
| CISD1 | Rabbit | Proteintech | 16006-1-AP | 1:5000 (WB) |
| Cone-arrestin | Rabbit | Chemicon | AB15282 | 1:250 (ICC) |
| COX4 | Mouse | GeneTex | GTX628901 | 1:1000 (WB) |
| CRALBP | Mouse | GeneTex | GTX15051 | 1:200 (ICC) |
| GFAP | Rabbit | Dako | Z0334 | 1:100 (ICC) |
| GS | Rabbit | Sigma-Aldrich | G2781 | 1:500 (ICC)  1:1000 (WB) |
| HO-1 | Rabbit | Enzo | BML-HC3001-0100 | 1:200 (ICC) |
| Iba-1 | Rabbit | WAKO | 019-19741 | 1:500 (ICC) |
| IL-33 | Rabbit | Abcam | Ab187060 | 1:250 (ICC) |
| Ki67 | Rat | Thermo Fisher | 14-5698-82 | 1:250 (ICC) |
| Kir4.1 | Rabbit | Proteintech | 12503-1-AP | 1:1000 (WB) |
| Lamin B1 | Rabbit | Abcam | Ab229025 | 1:1000 (ICC, WB) |
| Nestin | Mouse | Abcam | Ab6142 | 1:250 (ICC) |
| Nestin | Mouse | Thermo Fisher | MA1-110 | 1:1000 (WB) |
| Neurofilament-H | Rabbit | Proteintech | 21471-1-AP | 1:500 (ICC) |
| NIX (BNIP3L) | Rabbit | Cell Signalling | 12396 | 1:1000 (WB) |
| Notch1 | Rabbit | Abcam | Ab52627 | 1:100 (ICC) |
| Parkin | Rabbit | Cell Signalling | 2132 | 1:1000 (WB) |
| Pax6 | Rabbit | Abcam | Ab195045 | 1:350 (ICC)  1:1000 (WB) |
| PINK1 | Mouse | Ascites | AM6406a | 1:1000 (WB) |
| PGC1a | Rabbit | Novus | NBP1-04676 | 1:1000 (WB) |
| PKC-α | Rabbit | GeneTex | GTX130453 | 1:1000 (WB)  1:500 (ICC) |
| pUbSer65 | Rabbit | Sigma-Aldrich | ABS1513-I | 1:250 (ICC) |
| Rhodopsin | Mouse | Sigma-Aldrich | MAB5356 | 1:500 (ICC) |
| S100A | Rabbit | Abcam | Ab197896 | 1:250 (ICC) |
| S100β | Rabbit | Abcam | Ab52642 | 1:200 (ICC) |
| Sox2 | Rabbit | Abcam | Ab92494 | 1:100 (ICC) |
| TFAM | Rabbit | Genetex | GTX112760 | 1:500 (WB) |
| TOMM20 | Rabbit | Sigma-Aldrich | HPA011562 | 1:1000 (WB) |
| TSPO (PBR) | Rabbit | Abcam | Ab109497 | 1:10000 (WB) |
| Vimentin | Rabbit | Abcam | Ab92547 | 1:500 (ICC) |
| Vinculin | Rabbit | Thermo Fisher | 700062 | 1:1000 (WB) |
